# Biodiversity Research and Innovation in Antarctica and the Southern Ocean

**DOI:** 10.1101/2020.05.03.074849

**Authors:** Paul Oldham, Jasmine Kindness

## Abstract

This article examines biodiversity research and innovation in Antarctica and the Southern Ocean based on a review of 150,401 scientific articles and 29,690 patent families for Antarctic species. The paper exploits the growing availability of open access databases, such as the Lens and Microsoft Academic Graph, along with taxonomic data from the Global Biodiversity Information Facility (GBIF) to explore the scientific and patent literature for the Antarctic at scale. The paper identifies the main contours of scientific research in Antarctica before exploring commercially oriented biodiversity research and development in the scientific literature and patent publications. The paper argues that biodiversity is not a free good and must be paid for. Ways forward in debates on commercial research and development in Antarctica can be found through increasing attention to the valuation of ecosystem services, new approaches to natural capital accounting and payment for ecosystem services that would bring the Antarctic, and the Antarctic Treaty System, into the wider fold of work on the economics of biodiversity. Economics based approaches can be criticised for reducing biodiversity to monetary exchange values at the expense of recognition of the wider values of biodiversity and its services. However, approaches grounded in the economics of biodiversity provide a transparent framework for approaching commercial activity in the Antarctic and introducing requirements for investments in the conservation of Antarctic biodiversity by those who seek to profit from it.

## Introduction

This article examines the scientific and patent landscape for biodiversity based research and innovation in Antarctica and the Southern Ocean. The article is based on a review of 150,401 scientific articles and 29,690 patent families that make reference to the Antarctic or Southern Ocean in the open access Lens database of scientific and patent literature.

The Antarctic region is an important focus of scientific research in the context of the biodiversity and climate change crisis [1]. The impacts of climate change on terrestrial and marine biodiversity may be both positive and negative, with particular concern emerging over non-native species in terrestrial Antarctica and environmental warming and ocean acidification in the marine environment [1]. Commercial activity in Antarctica includes tourism and the harvesting of marine genetic resources such as Antarctic krill and Antarctic toothfish [2–4]. The region has also been a focus for bioprospecting or research on the potentially useful properties of Antarctic biodiversity for the development of new and useful products [5–10]. The emergence of commercially oriented research and development has led to increased debates around the governance of research activity, ethics and benefit-sharing. Debates on the governance of research and benefit-sharing mirror debates on access to genetic resources and benefit-sharing under the United Nations Convention on Biological Diversity and its Nagoya Protocol, and related policy processes such as negotiations on a new treaty on marine biodiversity in areas beyond national jurisdiction under the United Nations Law of the Sea. From 2005 onwards bioprospecting has appeared on the agenda of the Antarctic Treaty Consultative Meeting (ATCM) of Contracting and Consultative Parties to the Antarctic Treaty System (ATS). The Antarctic Treaty System consists of a set of agreements that aim to ensure that the Antarctic is a “natural reserve, devoted to peace and science” for the benefit of human kind. However, to date, activity under the Antarctic Treaty System with respect to bioprospecting has been limited to information gathering by the Scientific Committee on Antarctic Research (SCAR).

The aim of this article is twofold. First, we improve the evidence base for debates on the governance of research in Antarctica and the Southern Ocean by making datasets of scientific and patent literature and taxonomic data about the Antarctic publicly available through the Open Science Framework. The datasets are intended to contribute to methodological development in areas such scientometrics and machine learning based approaches to natural language processing [11–13,13–16].^2^ We argue that further methodological development is desirable, including by data providers, in order to address weaknesses in data coverage and data quality.

Second, we examine the main features of the scientific and patent landscapes for Antarctica and the Southern Ocean with a focus on biodiversity based innovation. The paper argues that efforts to address commercial research and development could usefully be approached in the wider context of the ecosystem services provided by Antarctic biodiversity [17–19]. This could be extended to the application of natural capital accounting, presently being incorporated into Systems of National Accounting (SNAs), to the Antarctic [20]. The rise of ecosystem services and natural capital accounting is grounded in increasing recognition within the economics community that biodiversity and the services it provides are not free and must be paid for. If we accept that biodiversity is not a free good and that everyone must, proportionate to their means, pay something we are able to ask other questions, such as: how much, by whom, in what form and to what ends? This paper does not aim to answer these questions but contributes to the evidence base for deliberation on the opportunities to address issues of fairness, equity and benefit-sharing for biodiversity based research and development in Antarctica and the Southern Ocean.

## Methods

This paper is a contribution from anthropology and data science that combines analysis of the scientific and patent literature with taxonomic data from the Global Biodiversity Information Facility (GBIF) on Antarctic biodiversity. The method consists of five main steps:

1. Capturing the raw universe of scientific and patent publications making reference to Antarctica and the Southern Ocean in multiple languages using the Lens open access database https://www.lens.org/;
2. Identifying and cleaning author, organisation, inventor and patent applicant names and linking with geospatial data sources using Microsoft Academic Graph (MAG) data tables (January 2019 release) from Microsoft Academic [21];^i^
3. Text mining the scientific literature and patent literature for taxonomic names with a focus on species names and a limited set of common names based on data from the Global Names Index (GNI) and GBIF; Refining the data to focus on scientific literature and patent data containing a verifiable Antarctic species using a cleaned version of Antarctic country code AQ data from GBIF; Text mining the results for Antarctic places names with a particular focus on patent data using data from the SCAR Composite Gazetteer of Antarctica (CGA) and the Geonames database of Antarctica (AQ) country code place names.

The steps above involved a number of elements and issues of interest to the data science community that can be summarised as follows.

Open access databases such as the Lens from Cambia and the Queensland University of Technology make it possible to search for data in multiple languages and to a more limited degree to search the full texts of scientific publications and patent documents. Based on a set of experimental tests the following multi-language query was developed to capture the available universe of publications about Antarctica and the Southern Ocean in multiple languages.

Antarctic* OR “Southern Ocean” OR “South Pole” OR “alqarat alqatabiat aljanubia” OR Antarctique OR Antarktida OR Antarktidë OR Antarktik OR Antarktika OR Antarktikí OR antarktis OR Antarktis OR Antarktisz OR Antarktyda OR Antartaice OR Antártica OR antártida OR Antártida OR Antàrtida OR Antartide OR Antartika OR Antartikako OR Jacaylku OR “Nam Cực” OR “namgeug daelyug” OR Suðurskautslandið OR Ανταρκτική OR Антарктида OR Антарктик OR Антарктика OR Антарктикийн OR Антарктикот OR Антарктыда

The Lens Scholarly database aggregates data from a number of different sources including Microsoft Academic Graph (MAG), Crossref (for meta data), PubMed for medically focused literature and CORE (core.ac.uk) for open access full texts. The available fields of search vary across data sources with all except for CORE being confined to metadata fields such as title, abstract, keywords, affiliations, authors etc. Table 1 summarises the raw search results from the different sources.

**Table 1:** Antarctic Paper Counts by Type

| name | papers |
| --- | --- |
| papers | 150401 |
| microsoft academic graph | 135150 |
| metadata | 122886 |
| CORE full texts only | 27515 |
| pubmed | 16053 |
| pubmed central | 2754 |

Note:
Metadata refers to Antarctic search terms in titles, abstracts, keywords, fields of study and MeSH terms.

In considering the raw data in Table 1 it is important to note two points. First, that the analysis in this paper is limited to the 135,150 papers from Microsoft Academic Graph. The reason for this is that the Lens does not directly provide access to affiliation data but it is possible to retrieve this data using the freely available Microsoft Academic Graph database tables. Second, cases where the Antarctic search query only appeared in CORE full texts merit more detailed investigation in future research. Except where they appear in Microsoft Academic Graph these texts are excluded from the quantitative analysis below.

The results of the search include any document that references Antarctica, the Southern Ocean or the South Pole anywhere in metadata (including author affiliations and bibliographic references) or the available full texts from CORE. This will inevitably include sources of objective noise, such as references to the South Pole of Mars or Titan or negations such as “except Antarctica”, and subjective noise such as the exploration of the role Antarctica plays in the human imagination in literary or cultural studies that may not be of interest to some readers. A conventional approach to dealing with noise in bibliometrics/scientometrics is to attempt to exclude it at source. However, we adopted a different approach informed by the possibilities of the rise of machine learning approaches to natural language processing and their future application to polar research.

Machine learning based approaches to Natural Language Processing (NLP) involve training models to engage in probabilistic classification of texts and named entity recognition (e.g. place names, species names). At the time of writing popular libraries include keras, fasttext, scikit-learn and spaCy (among others). The key condition for training models is the availability of preferably large volumes of labelled texts for use in training, testing and evaluating models. Viewed from this perspective, raw data that includes noise that is close to the subject matter (e.g. the South Pole of Titan or “everywhere except Antarctica”) is valuable. Rather than excluding noise at source we therefore adopted the approach of leaving the data as is and adding logical TRUE/FALSE columns to the raw data table as labelled filters. The filters are based on text mining of publication metadata (titles, abstracts, author keywords, fields of study, MeSH (medical subject heading terms). Table 2 displays the filters.

**Table 2:** Paper Counts by Subject (metadata only)

| name | papers |
| --- | --- |
| climate | 37015 |
| taxonomic name | 25662 |
| biodiversity | 25233 |
| southern ocean | 12939 |
| antarctic species | 12768 |
| arctic | 10965 |
| mammal | 8365 |
| planets | 4579 |

Table 2: Paper Counts by Subject (metadata only)
| name | papers |
| --- | --- |
| birds | 3758 |
| candida antarctica | 3751 |
| krill | 3127 |
| seal | 2848 |
| penguin | 2559 |
| whale | 1688 |
| acidification | 652 |
| innovation | 195 |
| ecosystem services | 122 |
| bioprospecting | 99 |

*Note:*
Counts of terms appearing in paper metadata including titles, abstracts, keywords, fields of study and MeSH terms.

The aim of the filters is to allow a user to restrict the data to areas of interest. For example, ‘taxonomic name’ is a filter for records containing a uninomial or binomial species name while ‘antarctic species’ refers to species that occur in Antarctica validated in the taxonomic data with an Antarctic location.

In the second step, data from the Lens was federated with Microsoft Academic Graph from Microsoft Academic (January 2019, release). Microsoft Academic Graph is based on data from the Bing search engine and is made available free of charge as a set of data tables that contain over 200 million scientific records. Federation was performed using a Databricks Apache Spark cluster on Microsoft Azure running R in RStudio with the *sparklyr* and *tidyverse* packages on the master node [21–24]. Data federation focused on table joins between the Lens data and affiliations and authors tables of Microsoft Academic Graph using the shared identifier (the paperid). This yielded an affiliation table with 5,021 identified organisations (affiliationid) and an authors table with 244,778 authors (authorid). One important and known limitation of Microsoft Academic Graph is that the affiliations data is incomplete [11]. Thus, 69,805 of the papers in the dataset were recorded with an affiliation id corresponding with 52% of the 135,150 papers. However, raw affiliation data is available in the authors table for the full MAG database. We used a multistep process described in the OSF repository to improve coverage to 99,794 (74%) of Microsoft Academic Graph data for Antarctica. The majority of the outstanding 34,249 papers were made up of book chapters, books and other data types that normally lack affiliation data (17,886). As a consequence, data on affiliations is incomplete and must be classified as indicative rather than definitive. While these results may give the scientometrics community reason for pause in using Microsoft Academic Graph, we would observe that interrogating these issues provides a basis for future improvements such as retrospective reindexing to pick up missing data.

With respect to patent data, at the time of the research the Lens included 115,915,955 patent documents from 63,366,633 families (publications grouped onto the earliest patent filing in a set) from 115 countries including regional and international patent offices. To retrieve patent data the same query was performed using full text search (titles, abstracts, description and claims). This yielded a raw count of 52,701 documents in 25,463 patent families from the search terms. The Lens is also important as a source of patent data for innovation research because it indexes scientific publications that are cited by patent documents. When these documents were added the total count of patent families rose to a raw 29,690 families.^3^

Patent documents are commonly republished multiple times. Thus, a single application may be republished as a patent grant or with an administrative search report or correction. The same application may also be submitted to multiple countries where it will also be republished. This introduces radical multiplier effects into patent counts. Thus, the 29,690 patent families in our raw set are linked to 163,615 later patent publications (family members). To control for this, patent analysts commonly reduce linked documents in a set or ‘family’ to the earliest first filing (known as the priority document). This article uses this approach. We added a “filing order” filter to the Lens patent data that reduces the original 29,690 Lens patent family documents to the 26,120 earliest first filings. Finally, it is important to emphasise that patent data, by virtue of access to the full text, is typically noisier than searches of the metadata for scientific literature with terms such as “South Pole” having multiple uses.

Text mining of the scientific and patent literature was performed in R using the *spacyr* package that provides access to the Python *spaCy* library for machine learning and Natural Language Processing and the R *tidytext* package [25,26]. Text mining focused on the identification of binomial and uninomial taxonomic names in texts followed by the identification of place names. This was performed by extracting noun phrases from the titles, abstracts, keywords, fields of study and MeSH terms for Lens records in the scientific literature. In the case of patent data, internal full text collections focusing on the US, the European Patent Office and the international Patent Cooperation Treaty were used to text mine the available titles, abstracts, descriptions and claims. To address memory issues when using *spaCy* with *spacyr* we used the *tidytext* package to parse texts into sentences, two word phrases (ngram 2) and words (ngram 1). It should be noted that approaches focusing on noun phrases are partly dependent on the language model (English) used for noun identification. We therefore expect room for improvement in data capture across multi-language sources.

Matching with taxonomic names and place names was performed using dictionary based approaches. Noun phrases were matched against a dictionary of just over 6 million binomial species names originally extracted from the Global Names Index (GNI) and its web service at http://gni.globalnames.org/ [27]. The full list of binomials was derived from a copy of the Global Names Index kindly provided by David Remsen and Dmitry Mozzherin as leading developers of the wider Global Names Architecture. Individual words (uninomials) were chosen for matching with entries in the Families of Living Organisms (FALO) dataset from GBIF that consists of single or uninomial names for Kingdoms, (e.g. Animalia), Families (e.g. Ursidae for the bear family) etc. [28,29]. A 2014 species list from the World Register of Marine Species (WoRMS) database was used to add a filter for marine species in the literature and patent data tables. We would note that careful attention is required to improvements in the classification of marine species (e.g. to distinguish between terrestrial aquatic and marine organisms) in later updates of WoRMS when approaching this filter.

The raw results of text mining with dictionaries were passed to the GBIF API using the *taxize* package from ROpenSci to retrieve the taxonomic hierarchy [30]. One issue when retrieving the taxonomic hierarchy for thousands of species is that a single species name may match to multiple records (e.g. as synonyms or homonyms). However, it is impractical to manually review thousands of results when retrieving data. Fortunately, the return from *taxize* includes a ‘multiple matches’ column that identifies these cases. The multiple matches filter is retained in the taxonomic data tables to allow taxonomic specialists to review and, as necessary, refine the data.

Scientific and patent publications that include taxonomic names commonly include multiple names. This is particularly true in patent documents and presents the challenge that a particular organism may or may not occur or have been collected in the Antarctic. GBIF maintains a dataset of occurrence records (observations) with country code AQ that in May 2019 consisted of 2,729,211 occurrence records [31]. However, at that time, over 1 million of the records were recorded at latitude -91 or -90 revealing unlikely and invalid records. To address this, the data was restricted to records containing a text entry for locality and a second data set for -60 latitude South was generated and combined [32]. To address noisy records a multi-step procedure was adopted involving removing inaccurate coordinates with the ROpenSci *CoordinateCleaner* package in R [33]. In the second step, the SCAR Composite Gazetteer of Antarctica (CGA) of 23,833 names, was used to text mine the locality field in GBIF data and single occurrence records were manually reviewed in VantagePoint from Search Technology Inc. In the third step, single species occurrence records that lacked locality information were identified. In the fourth step, a filter was added for occurrences south of -60 degrees latitude as the demarcation point for the Southern Ocean and Antarctica. In the fifth step, a species occurrence count was added based on the observation that low species occurrence records that lack locality information are often noise. In a sixth step, a filter was added for fossil records based on the existing GBIF “basis of record” field. Occurrence records with a validated Antarctic location in the locality field became the basis for the ‘antarctic species’ filter applied across the dataset. The addition of an ‘occurrence count’ field allow the species related data to be progressively restricted to those with a validated Antarctic location in an ordered way.

As this summary of methodological steps makes clear, the federation of scientific literature, patent literature and taxonomic data involves a number of methodological challenges. It is also clear that while the rise of open access databases revolutionises the opportunities for this type of analysis at scale, there are a variety of limitations in the data sources. This means that the analysis presented in this paper is indicative rather than definitive. Nevertheless, highlighting these limitations presents opportunities to identify ways forward in improving data coverage and data quality to inform decision-making.

## Results

Figure 1 displays an overview of the raw dataset for the Antarctic search terms. In Figure 1A we can immediately observe that after a steep increase in the paper count to a peak in 2014 of 7,468 publications the data displays a declining trend. However, in our view this will reflect data availability issues with Microsoft Academic Graph rather than an actual decline in publications referencing Antarctica. The reason for this is that a steep decline from around the same point is observable for non-Antarctic data. An explanation of this issue could usefully be added to the Microsoft Academic Graph documentation to improve certainty for users.

**Figure 1:**
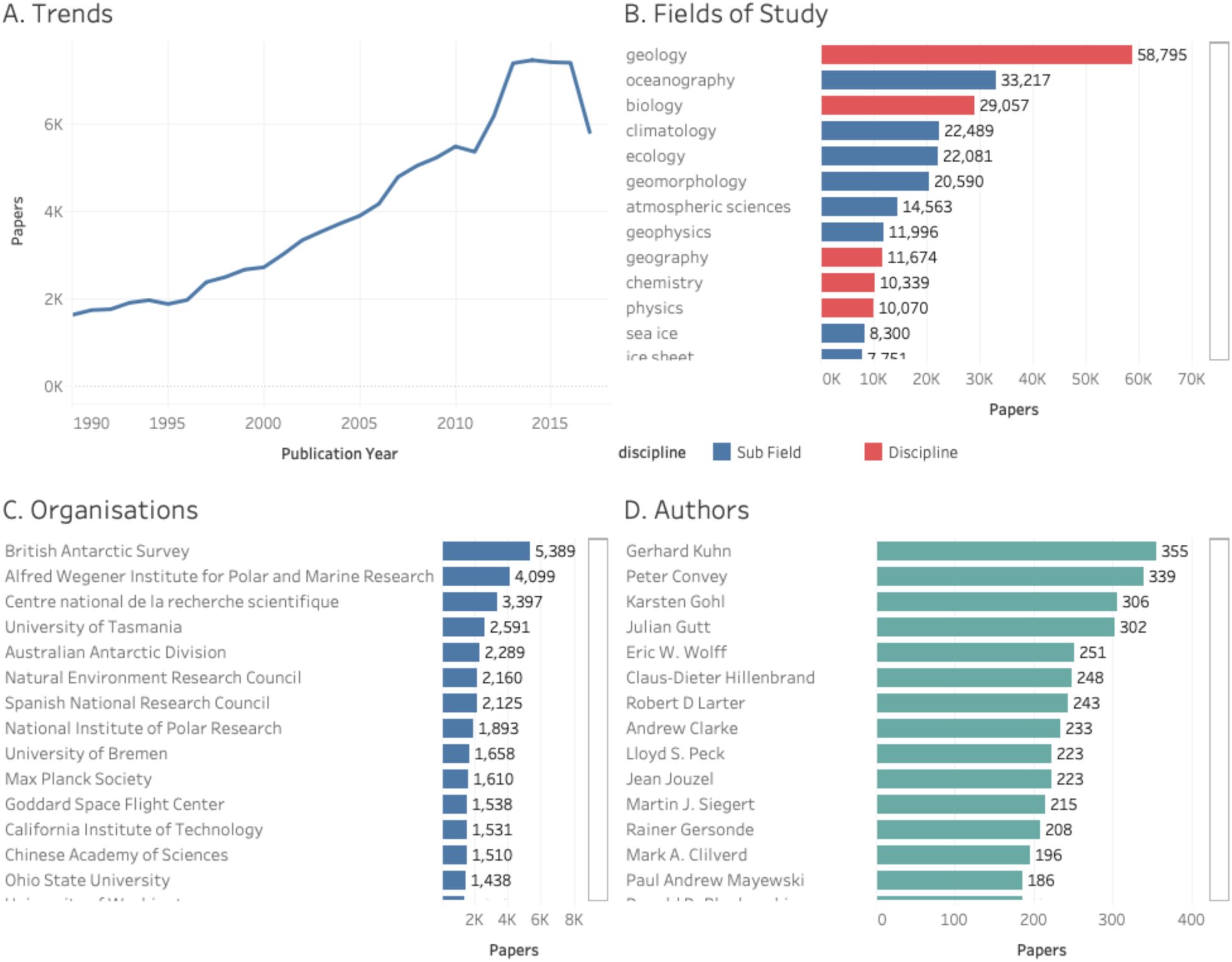
Overview of Scientific Literature for Antarctica.

Microsoft Academic Graph uses a combination of data from Wikipedia and machine learning to identify and label papers by subjects called “Fields of Study” [34]. In contrast with approaches such as Clarivate Analytics Web of Science, that categorise journals rather than papers, this approach allows for the use of multiple labels at different levels of detail [34].

In the January 2019 release, MAG Fields of Study consisted of 19 top level disciplines that are displayed in red in Figure 1B. The remaining fields, shown in blue, are children of the MAG disciplines. Thus, in Figure 1B oceanography, climatology, geomorphology, atmospheric sciences etc. are all children of geology. In contrast, ecology and botany are children of biology. These children in turn have sub-child labels at varying levels of detail including limited labels for taxonomic classification. Overall, this signifies that papers may be divided into very broad fields and may appear multiple times in the rankings at different levels of detail.

Figure 1C displays the available data on the number of papers per organisation. The data is counted by aggregating the papers linked to an organisation (which may include multiple authors from the same entity) and then counting the distinct papers. As noted above, it should be emphasised that this data is indicative rather than definitive. As the resolution of affiliation data improves we would expect the numbers and relative positions of organisations in the rankings to change. Nevertheless, the data is indicative of some of the most important organisations conducting research involving the Antarctic in recent decades.

Researchers from 134 countries appeared in the raw publication data relating to the Antarctic. However, rankings are affected by the availability of affiliation data. We can gain an initial idea of the geographic distribution of organisations involved by mapping organisations in the data that also appear in the public domain *Global Research Identifier Database (GRID)* https://www.grid.ac/. The GRID database forms part of a growing effort to harmonise institutional names for geographic mapping and other purposes. Figure 2 breaks out the full data from Figure 1C and displays a map of available geographic data for organisations publishing research relating to Antarctica and is accompanied by a ranking of countries based on the number of distinct publications of all types linked to Antarctica.

**Figure 2:**
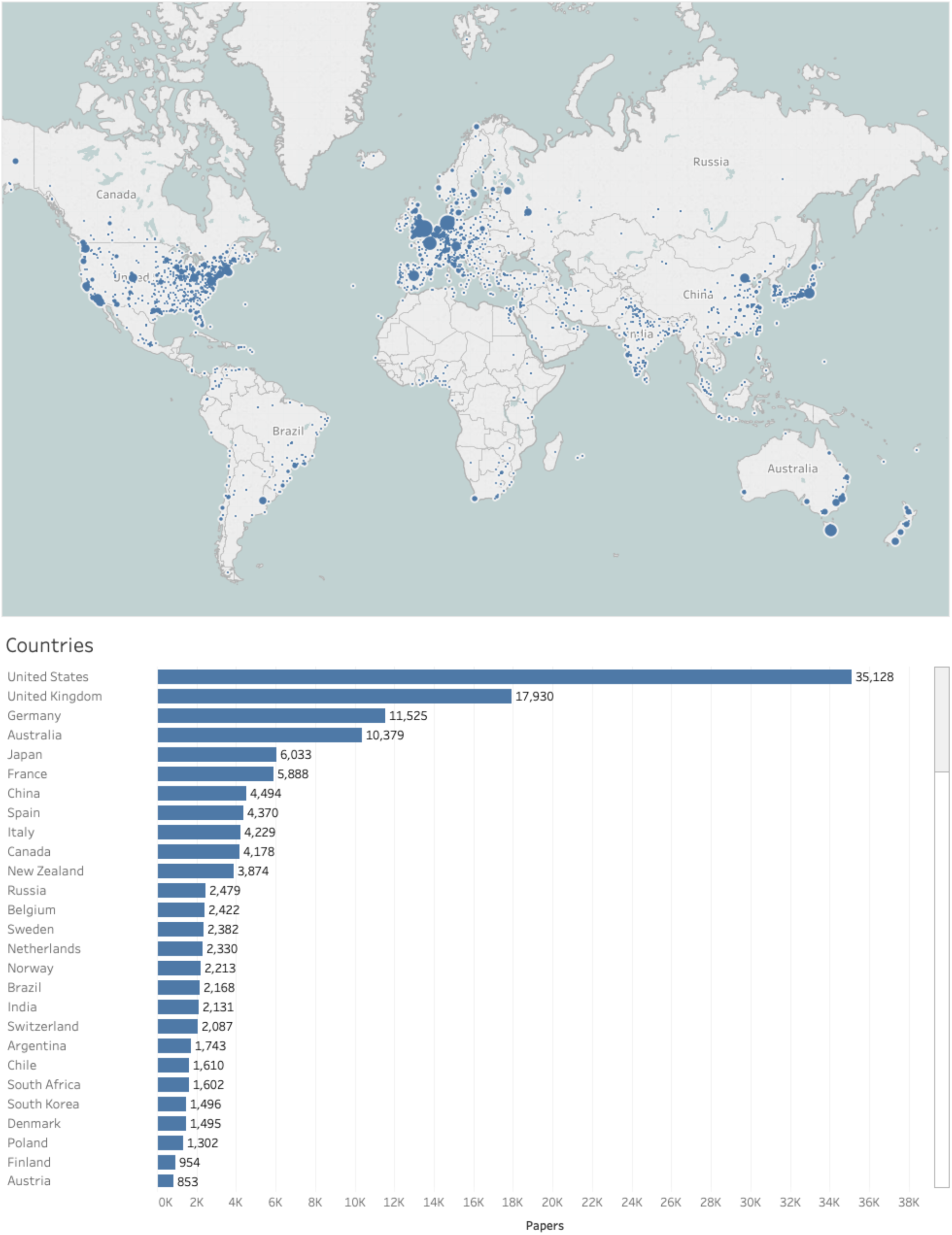
Geographic Distribution of Research Organisations Linked to Antarctica.

It is worth noting that some countries with organisations with a significant presence in Antarctic research are probably under-represented in the organisation map because their data is distributed across multiple organisations with no available georeference data, notably Russia (with 63 organisations).

Figure 1D (above) displays the rankings of papers by individual authors. Top ranking authors, based purely on the number of published papers or datasets appearing in Microsoft Academic Graph, include marine geologist Gerhard Kuhn at the Alfred Wegener Institute Helmholtz Centre for Polar and Marine Research [35], terrestrial ecologist Peter Convey at the British Antarctic Survey [3,36], and geophysicist Karsten Gohl at the Wegener Institute [37,38]. Leading women scientists in the data by publication count include geophysicist Gabriele Uenzelmann-Neben at the Wegener Institute [39,40], marine biologist Katrin Linse at the British Antarctic Survey [41,42] and climate scientist Valerie Masson-Delmotte [43,44]. In some cases researchers may be active in research and publication on both Antarctica and the Arctic as part of wider polar research.

This global overview of research referencing the Antarctic serves to demonstrate the potential of tools such as the Lens and Microsoft Academic Graph to illuminate research landscapes on the global level. At the same time, data on trends, affiliations and georeferencing exposes the need for improvements in data quality and coverage. However, while recognising these constraints, this approach also significantly expands our access to data on scientific publications about the Antarctic. In an important contribution to bibliometric analysis Ji et al. 2014 published analysis of research on publications in the Antarctic between 1993 and 2012 using a search for the Antarctic in Web of Science that yielded 36,238 publications (after the exclusion of species containing antarctica in the name) [45]. In contrast, for the same period Microsoft Academic Graph produced 71,804 distinct papers with 79,647 across the Lens. The increase in publication data will reflect a combination of the choice of search terms, the wider scope of Microsoft Academic Graph, the growing availability of data in multiple languages (with 46 languages represented in the data), the growing availability of millions of open access full texts through CORE (core.ac.uk), and the growing emphasis on open access data in scientific policies.

The increasing availability of publication data at scale brings with it a need to focus on potential sources of noise but also provides opportunities to drill into the data in specific areas of interest. Existing bibliometric research on the Antarctic has focused on the exploration of highly cited research [46], the role of research stations in promoting collaborative research [47], and mapping glacier research with Web of Science [48]. As this suggests, publication data on Antarctica provides rich opportunities for the exploration of specific research themes. We now turn to the analysis of research on Antarctica involving biodiversity at the species level as a basis for exploring commercial interest in Antarctic species in patent data.

### Biodiversity Research in Context

As we observed above, the research profile for the Antarctic is dominated by geology, climatology and other Earth Science subjects. This is reflected in the top cited publications for Antarctica including topics such as: high resolution interpolated climate surfaces for global land areas [49], mixed effects modelling of ecology with R [50], the IPPC 4 report on *Climate Change 2007: Impacts, Adaptation, and Vulnerability*, global analysis of sea surface temperatures [51], and the climate and atmospheric history of the past 420,000 years from the Vostok ice core, Antarctica [52].

As we observed in the discussion of Antarctic fields of study, biology is a prominent subject area that is accompanied by a number of large subfields such as ecology, botany and biochemistry. Top cited research in the field of biology includes a new phylogenetic method for comparing microbial communities that includes comparison of Antarctic and Arctic communities [53], the influence of temperature on phytoplankton growth [54], sterol markers for marine and terrigenous organic matter [55], analysis of the genus *Nocardiopsis*, including discussion of *Nocardiopsis antarctica*, as a distinct Actinomycete lineage [56], and fatty acid trophic markers in the pelagic marine environment [57].

The main focus of the present research was on identifying and extracting species level information from research on the Antarctic using text mining. As a starting point, research on species can be divided into two broad categories: a) direct field research involving Antarctic species, and; b) indirect or follow on research, including classification and comparative analysis, and the exploration of the properties of organisms.

In total we identified 1,819 binomial species names with recorded occurrences in the scientific literature for the Antarctic. Of these, 1,666 had specific locality information. In the case of some animals such as whales, seals, penguins, and krill, common names, e.g. Blue whale or Adelie penguin, appear more frequently in the literature than their Latin names. To address this, additional counts were performed for the major groups including both common and taxonomic names and marked in the accompanying data table. Information on a public collection of biodiversity literature for Antarctica and the Southern Ocean is provided in the supplementary material.

Figure 3 displays the data ranked by species and the number of scientific publications for the 1,819 species.

**Figure 3:**
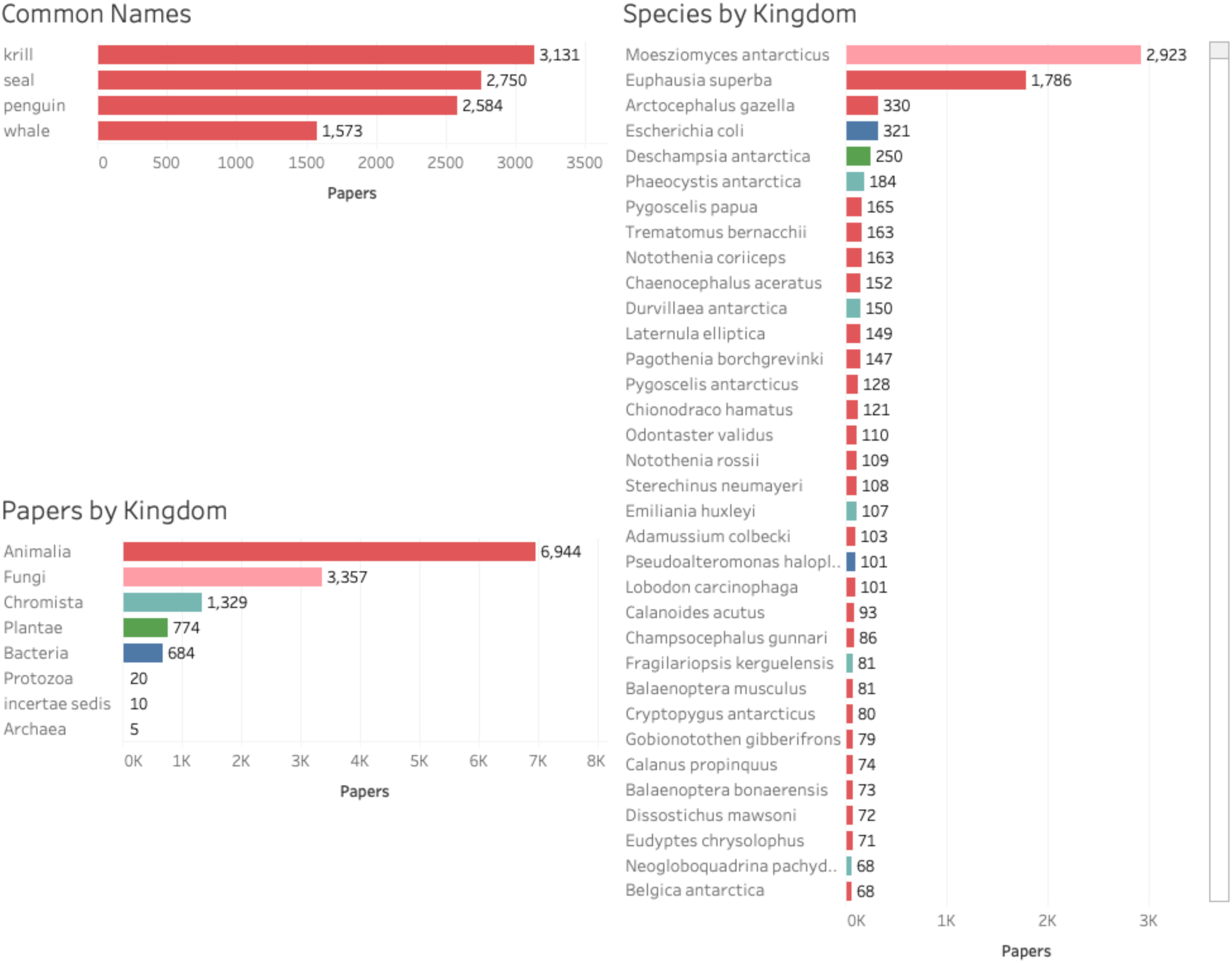
Top Ranking Species for Antarctica.

The data in Figure 3 reveals the prominence of *Candida antarctica* (accepted name *Moesziomyces antarcticus*) and krill (*Euphausia superba*) outside the major Antarctic mammals. This reflects the economic importance of *Candida antarctica* and the ecological and economic importance of krill.

We gain a more detailed insight into the prominence of species across the major kingdoms in the scientific literature in Figure 4. We will now briefly summarise some of the highlights of the literature and begin to focus in on research with commercial applications.

**Figure 4:**
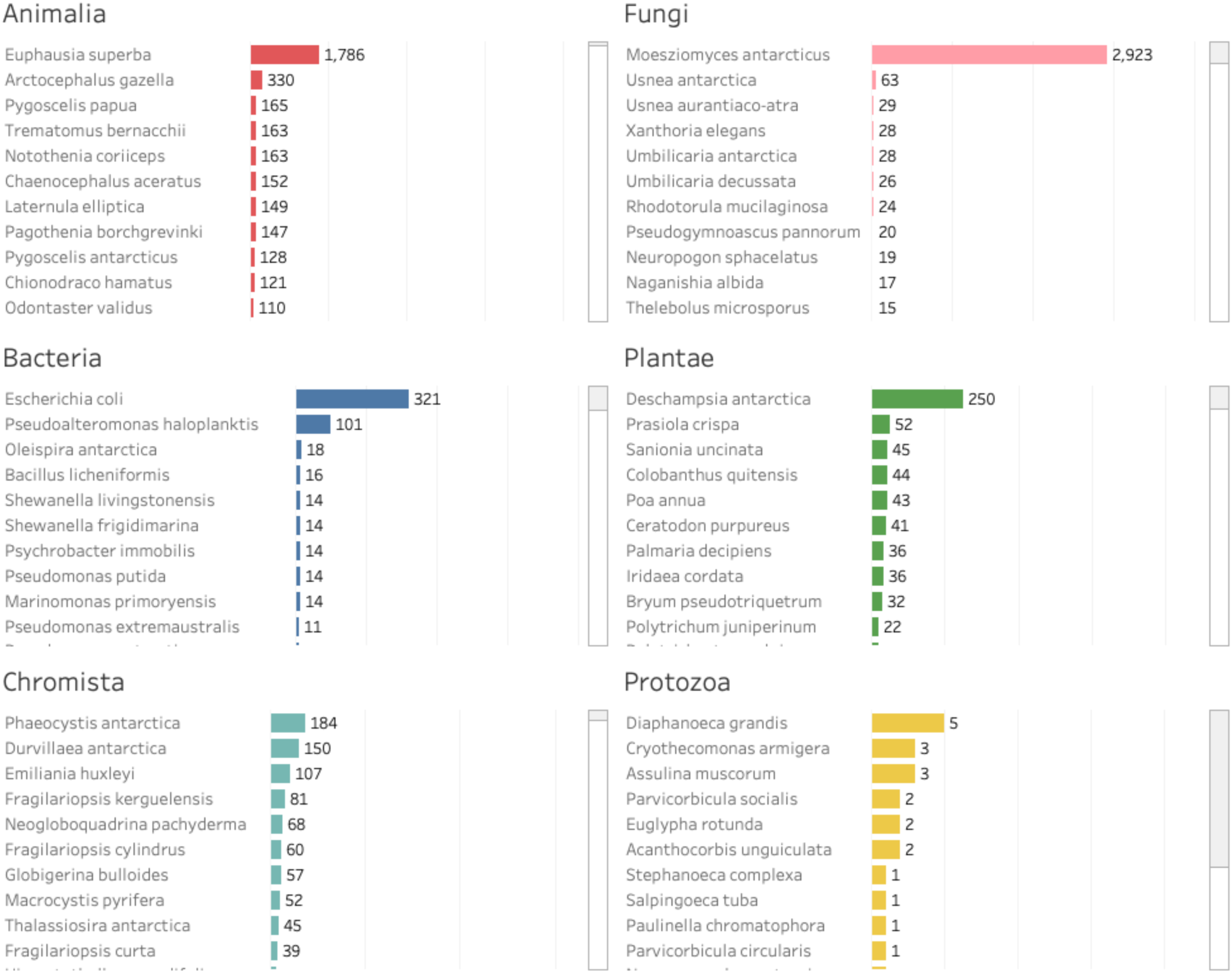
Top Ranking Species for Antarctica by Kingdom.

For animals on both common and taxonomic names Antarctic krill is a major focus of the literature that reflects significant economic interest in oil extraction from an abundant species that is rich in omega-3 polyunsaturated fatty acids [58–60]. Much of the literature on Antarctic krill details methods and success-rates in extracting proteins, fatty acids, amino acids and lipids from this species. Additional work has focused on the suitability of krill species as feed in salmon aquaculture [61,62]. The combination of climate change and commercial exploitation has led to work to model the impacts of any decline in Antarctic krill biomass on predators [63]. Antarctic krill are also a focus of the ecosystem-based fisheries management approach of the Commission for the Conservation of Antarctic Marine Living Resources (CCAMLR) [64]. Recent work on krill reveals concern that while ecosystem services in the Southern Ocean may increase under climate change this may occur at the expense of the decoupling of ecosystem provisioning for endemic species [65]. In other words, the food supply for endemic species may be disrupted leading to a need for specific management of biodiversity [65].

The Antarctic fur seal, *Arctocephalus gazella*, a historic focus of the seal fur trade, has been a focus of basic research on foraging behaviour and diet with recent research examining the impact of human associated *Escherichia coli* in pinnipeds such as *A. gazella* [66–68].

In contrast, the Emerald rockcod (*Trematomus bernacchii*) is a focus of interest because it has lost the ability to produce heat shock proteins in response to thermal stress, a capacity once regarded as universal amongst organisms [69,70]. In total we identified 382 articles for Antarctic fish including notothenioids, such as *Notothenia coriiceps* and members of *Trematomus*, with cold adaptation as a significant focus of research (see below).

In the case of Fungi the data is dominated by *Candida antarctica*. *Candida antarctica* and *Pseudozyma antarctica* are synonyms for the accepted name *Moesziomyces antarcticus*. As the literature is dominated by the use of the synonym *Candida antarctica* we will continue with that practice. The type specimen for *Candida antarctica* was originally collected from sediment at 9 metres depth from Lake Vanda in Victoria Land, Antarctica (mycobank specimen record 19800). Lipases from this species have been used for a wide range of purposes. B-component lipase derived from this yeast has been found to be a significantly robust lipase. It is highly stereospecific, and has been used as a biocatalyst in a wide array of chemical reactions, with uses in biotechnology, bioengineering, biochemistry and biofuels.[71–75] *Candida antarctica* lipase B has also been found to be highly effective in aiding the dissolution of carbohydrates [77] and for the production of amines and amides. As Gotor Fernandez et. al. 2006 explain “Simplicity of use, low cost, commercial availability and recycling possibility make this lipase an ideal tool for the synthesis and resolution of a wide range of nitrogenated compounds that can be used for the production of pharmaceuticals and interesting manufactures in the industrial sector.” [78].

The most prominent bacteria in the Antarctic literature is the ubiquitous *Escherichia coli* or *E. coli*. The prominence of *E. coli* in the Antarctic literature mainly arises from its use as a research tool [79–81]. Examples of the use of *E. coli* in Antarctic research include a dosimeter to evaluate the penetration of biologically active ultraviolet radiation within a water column based on the sensitivity of a particular strain of *E. coli* to ultraviolet radiation [82]. However, *E. coli* also appears in the taxonomic record for Antarctica through records from Davis Station. Growing interest in the implications of the increasing presence of humans in Antarctica are reflected in exploration of the impacts of *E. coli* strains in human waste upon the Antarctic environment [68,83]. As noted above, the impacts of human associated *E. coli* have also become a focus of research in seal populations [68]. Research on the health of penguin populations has identified antibiotic resistant bacteria such as *E. coli* in Gentoo penguin breeding areas [84]. The discovery of antibiotic resistant strains of *E. coli* in penguin populations implies that human activities are responsible [84].

*Pseudoalteromonas haloplanktis*, recorded in the taxonomic record at Frei Montalva Base on King George Island, appears in over 100 publications. The majority of research tends to focus on the capacity of this species to exist at cold temperatures [85–87]. Beta-galactosidase from this species has been shown to outperform other commercial beta-galactosidases suggesting that the cold-adapted beta-galactosidase could be used to hydrolyse lactose in dairy products processed in refrigerated plants [88]. The bacterium *Oleispira antarctica*, recorded in the taxonomic record at Road Bay in the Ross Sea, appears in 18 publications, two of which have received 150 or more citations suggesting significant interest. This interest appears to arise from *O. antarctica’s* hydrocarbon degrading properties which may be beneficial in the bioremediation of oil spills [89,90].

In the case of plants, Parnikoza et al. 2011 highlight that *Deschampsia antarctica* and *Colobanthus quitensis* are the only two flowering plants that have colonized the Maritime Antarctic [91]. As we will see below they are of significant commercial interest in connection with cold resistance [92]. *Deschampsia antarctica* appears in 250 papers and dominates the data on plants. Existing research suggests that *Deschampsia antarctica* may be useful as a bioindicator of climate change in Western Antarctica [93] while more recent work focuses on the mechanisms that allow it to survive in the Antarctic environment [94] and how these mechanisms fare in conditions of warming temperatures [95–97]. Difficulties in the interpretation of the taxonomic record for Antarctica are reflected in the presence of the Australian seagrass *Amphibolis antarctica* in the raw taxonomic data which as far as we can establish, despite its name, does not have a recorded distribution in Antarctica or the Southern Ocean.

For chromists, single and multicellular eukaryotes including some algae and diatoms, scientific attention has focused on *Phaeocystis antarctica* and *Durvillaea antarctica* (New Zealand Bull Kelp). The marine phytoplankton *Phaeocystis antarctica* has been a focus of analysis in connection with the formation of algal blooms and carbon sequestration in the Southern Ocean and their role in the carbon cycle [98]. Recent work has focused on issues such as the role of iron in colony formation and the impacts of iron limitation and ocean acidification on *P. antarctica* [99,100]. This in turn is linked with wider research on the implications of ocean acidification for diatoms and other marine organisms in Antarctica [101]. Research on *P. antarctica* also involved simulation of iron fertilization that can be linked to models for geoengineering experiments [102,103].

*Durvillaea antarctica* appears to be quite widely distributed in the Southern Ocean and countries such as New Zealand and Chile. Research on this species includes work on kelp rafts in the Southern Ocean and subantarctic, including the role of kelp mats in the dispersal of marine bivalves [104,105]. More commercially oriented research is reflected in work on the nutritional content of the edible *D. Antarctica* [106]. The anaerobic digestion of this species to produce biogas has also been evaluated as a method for producing renewable energy [107]. Research has also been conducted to extract soluble β-1,3/1,6-D-glucan from this species, which has been indicated to have immunostimulant properties [108]. High-M alginate extracted from this species, which has also been shown to have immunostimulatory properties, has been used in studies to create a dietary supplement for feeding and weaning Atlantic cod [109].

Protozoans have received relatively little scientific attention in Antarctic research to date. *Diaphanoeca grandis* isolated from saline Antarctic lakes and coastal sites in research dating to the early 1990s has received the greatest attention so far [110–112]. Research on *Bicosta spinifera* dating to the early 1980s is also limited but has focused on issues such as seasonal variation in abundance [113] with more recent work reporting on the Polarstern project in the Weddell Sea [114]. In recent work, *Cryothecomonas armigera* is being used in work to develop a bioassay to inform water specific guidelines to address pollution in Antarctica [115–117].

Research on Archaea, single celled microorganisms, in Antarctica appears to be very limited (not shown in Figure 4). The majority of research has focused on *Methanococcoides burtonii*, with over 30 publications. However, a number of these papers have been relatively highly cited such as work on genomics, proteomics and membrane lipid analysis in understanding mechanisms for cold adaptation [118–120]. Additional work has also taken place on *Halorubrum lacusprofundi* focusing on amino acid substitutions in cold adapted proteins [121].

Viruses have very limited coverage in GBIF data and are therefore not picked up in text mining with this data source. However, as we might expect, research on viruses appears in 472 publications for other species in the data. This includes viruses in Antarctic lakes [122], research on viruses and antibodies in Antarctic seals [123,124], a wider review of research on viruses in cetaceans [125,126] and research on viruses in penguin populations [127,128].

Our purpose in this section has been to provide a brief overview of biodiversity research in Antarctica and the Southern Ocean and to begin to focus on research activity with actual or potential commercial value. We turn now to the growing body of literature on bioprospecting, or biological research with a commercial focus, in the Antarctic.

### The Bioprospecting Literature

A significant literature has emerged that makes reference to bioprospecting or biological prospecting in Antarctica, consisting of over 90 articles. These articles range from research with a specific focus on identifying the potentially useful properties of Antarctic organisms to consideration of the policy implications of commercially focused research and development for the Antarctic environment and benefit-sharing.

The most highly cited article on bioprospecting in the Antarctic is a 2013 analysis of fungal communities associated with macroalgae in Antarctica with potential bioactive compounds that has so far received 83 citations [129,130]. Other research is comparative in nature, such as comparing samples of soil bacteria from arid Brazilian and Antarctic soils that are capable of digesting cellulose [131,132]. Still other work focuses on methodological development such as improved culturing from metagenomic (environmental) samples from cold environments [133,134]. Innovation in research methods for bioprospecting research also extends to the use of genome editing techniques and single cell sequencing for organisms from terrestrial and marine ecosystems [135,135,136]. Work on methods and techniques frequently refers to polar regions rather than necessarily involving direct field research. This is also reflected in review articles on issues such as fungi from terrestrial and marine Antarctic environments [137]. Recent literature on bioprospecting that has yet to attract significant citations includes work on enzymes from filamentous fungi [138], Antarctic bacteria as a source of novel antibiotics [10], and as sources of antimicrobial, antiparasitic and anticancer agents [139,140]. We would emphasise that the literature using the term bioprospecting has not increased dramatically over the years from the first record in 2002, with a peak of 8 publications in the available data for 2018 and an average of 4 publications a year between 2002 and 2018. In our view the use of the term bioprospecting will prove to be an unreliable indicator for what we would prefer to call commercially oriented research and development focusing on the potential applications of the properties of Antarctic organisms.

Bioprospecting also became an increasing focus of policy research from the early 2000s onwards in connection with potential measures under the Antarctic Treaty System (ATS) and is situated in a wider emerging literature on the governance of areas beyond national jurisdiction [141,142]. This includes potential legal and policy measures [6,143]. The ethics of commercial exploitation of Antarctica and Southern Ocean resources has also recently emerged as an important topic notably in a 2020 special issue of *Ethics in Science and Environmental Politics* [144–147].

Debates about bioprospecting in Antarctica have been closely tied up with patent activity. In the economics literature patent activity is used as a proxy output indicator for otherwise invisible investments in research and development [6,7,148]. That is, the filing of a patent application is an outcome of underlying financial investments in research and development [6,7,148]. In contrast, in wider policy debates on biodiversity, the filing of a biodiversity based patent application has become associated with the concept of biopiracy, or misappropriation, of genetic resources from countries and communities for commercial gain without returning benefits to countries, communities or biodiversity conservation. We now turn to the available data on patent activity for biodiversity from the Antarctic.

### Patent Activity

We identified patent activity referencing Antarctica using the search strategy described above across the full texts of patent documents worldwide. The raw data was reduced to 29,690 applications and then further reduced to 26,120 earliest first filings that form the basis of patent families. We then text mined the documents for any type of species name and reduced the results to those with a verifiable occurrence in Antarctica or the Southern Ocean in the available taxonomic record from GBIF. We identified a total of 3,907 patent applications and 2,738 first filings that contained a verifiable Antarctic species. In total we identified 1,212 species in the patent data of which 354 were verifiable Antarctic species based on locality information in the taxonomic record.

In approaching this data we would note that the data on Antarctic species that formed the basis for the search will inevitably be incomplete. As discussed below, we also note that the appearance of an Antarctic species in a patent document does not necessarily mean that an element of that species is claimed by the applicants. We will begin with an overview of the patent data containing Antarctic species and then progressively narrow the focus before concluding with examples of direct collection of samples in Antarctica.

Figure 5 displays the counts of species appearing in the full texts of patent documents that are known to occur in Antarctica.

**Figure 5:**
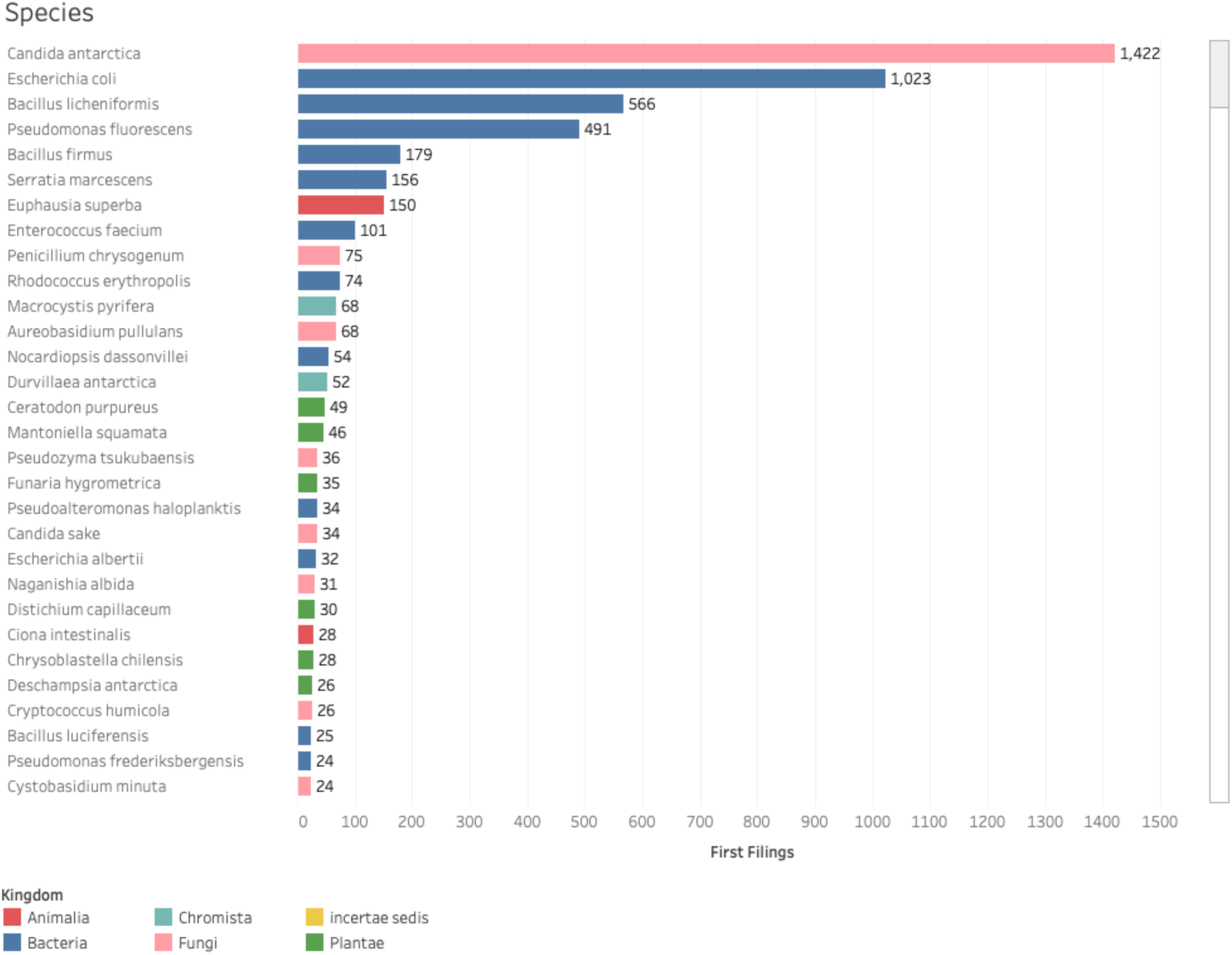
Antarctic Species in Patent documents Ranked by First Filings.

Figure 5 reveals that, as we might expect from the scientific literature, the top species is *Candida antarctica* (accepted name *Moesziomyces antarcticus*). This is followed by the ubiquitous *E.coli*. The presence of widespread species such as *E. coli* will in our view reflect the use of this organism as a tool in biotechnology rather than specific strains from Antarctica. This will also be true for other widely distributed species that have been recorded in the Antarctic.

One important feature of patent activity is that a species may be mentioned in different sections of a document. As a general rule, patent documents that mention a species in the title, abstract or claims will in some fundamental sense involve that species in the invention, either as a source for the invention, such as a lipase, or as a target of the invention such as a pathogen. However, the main density of species references is found in the description section. Figure 6 shows the breakdown of species names in the patent data presented in Figure 5 by document section ranked on patent claims.^4^

**Figure 6:**
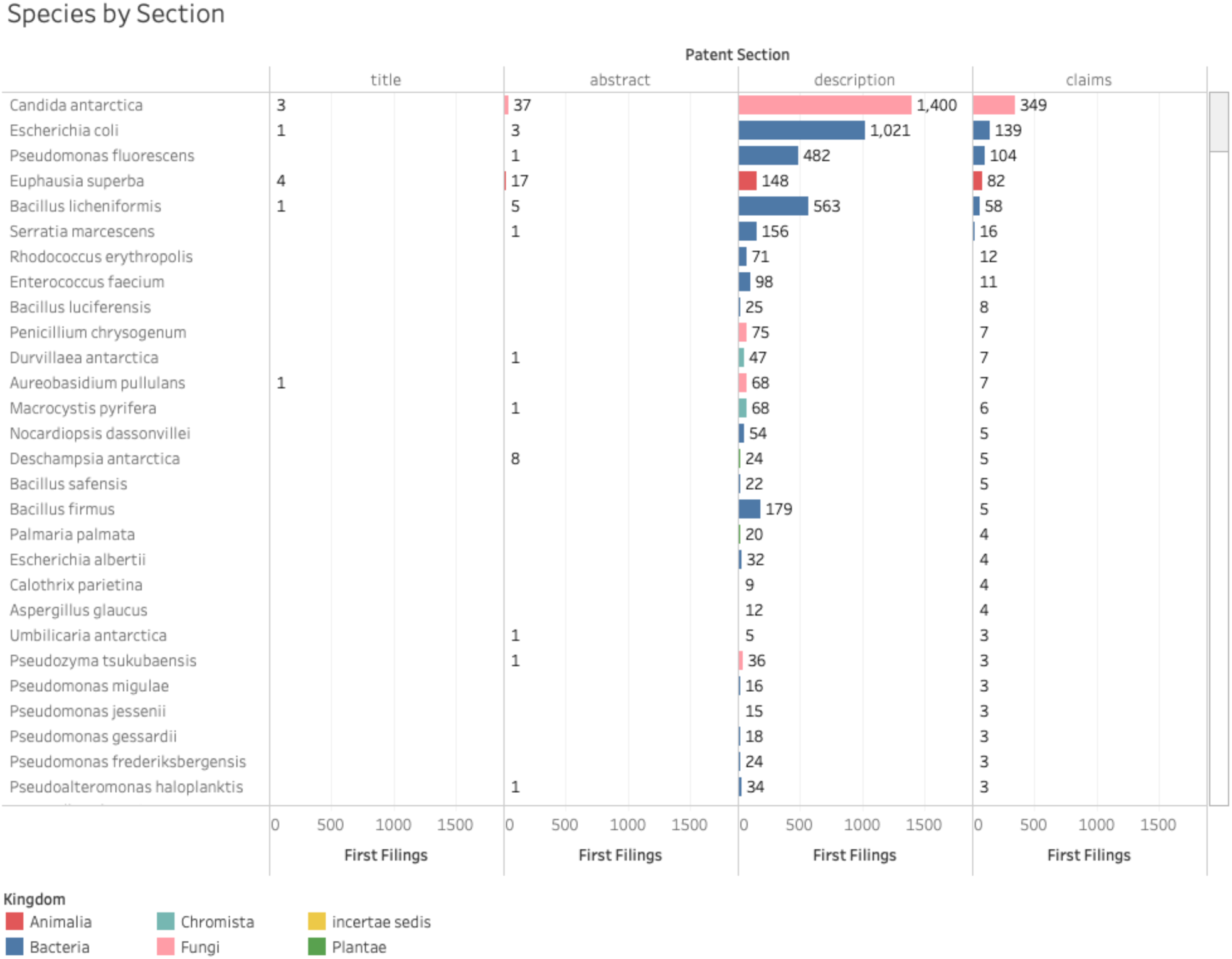
Antarctic Species in Patent Documents by Section Ranked on Claims.

As Figure 6 reveals the majority of references to a species appear in the description section with the remainder appearing in the claims.

References to species may appear in an application for a number of different reasons:

- As part of the claimed invention (the species is material to the invention);
- As part of experiments leading to the claimed invention;
- As an actual or potential component or ingredient in the invention, including in claims constructed on the genus, family, phylum or higher taxonomic levels;
- Literature citations (see below);
- Passing references (e.g. “in every species except…”, or “species x has been used to do y”) and long lists (notably for viruses);
- As DNA or amino acid sequences that are either used as comparative reference sequences or claimed.

In practice, determining whether a species is material to a claimed invention requires close attention to and interpretation of the texts. In the discussion below we provide examples of the different reasons that a species may appear in the text. Figure 7 presents an overview of the 2,738 first filings.

**Figure 7:**
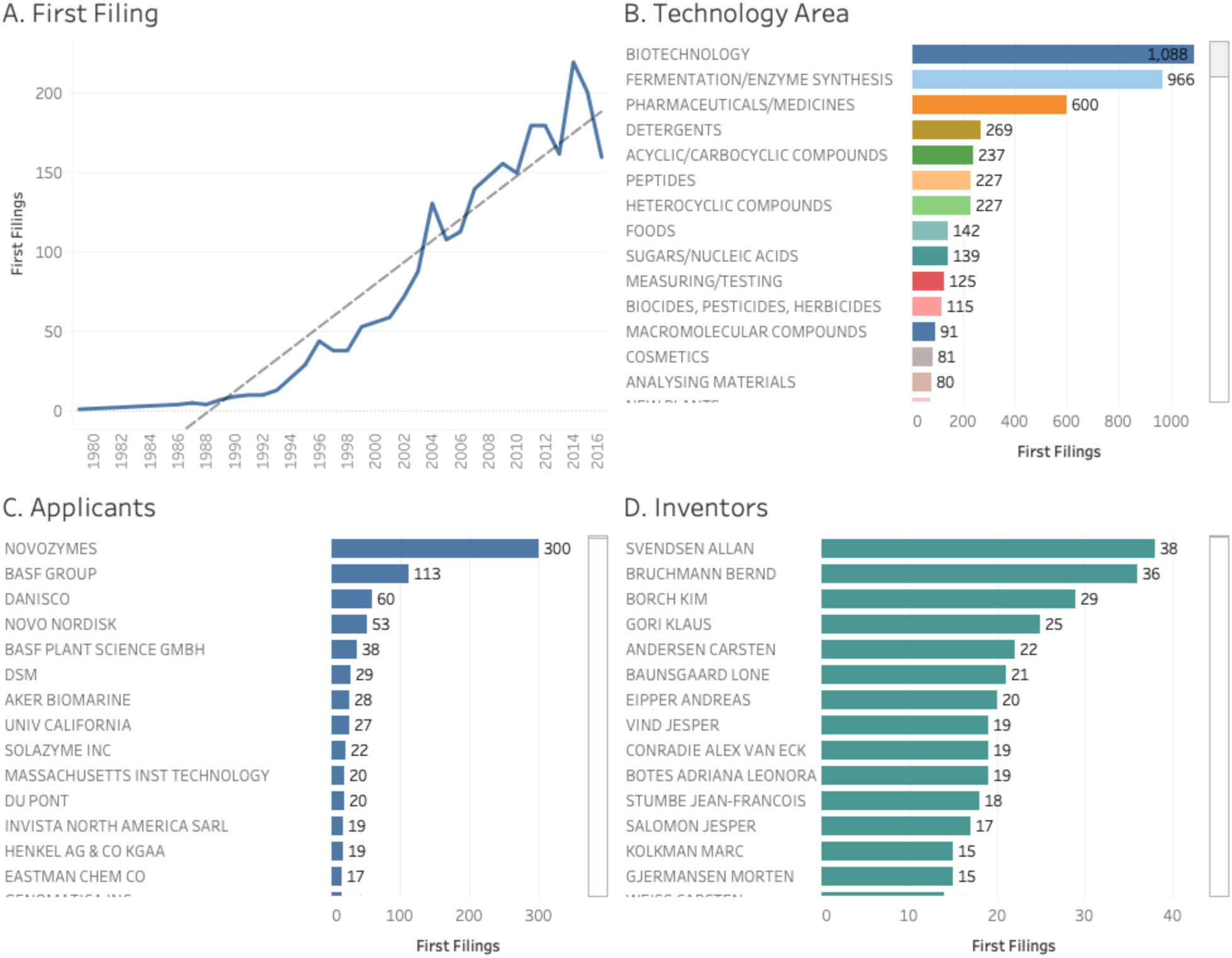
Overview of Patent Activity involving Antarctic Organisms.

Figure 5A reveals a rising, if irregular trend in filings. The apparent decline in filings in 2016 will normally reflect a data lag time of at least two years between the filing of a patent application and its publication. While a rising trend is observable from 2000 onwards the overall number of filings, peaking at 220 in 2014, is relatively modest, particularly when outstanding issues such as noise in the form of species recorded in the Antarctic that were not collected in the Antarctic are taken into consideration.

Figure 5B presents data on the main technology areas based on International Patent Classification subclasses and has been edited for readability. Figure 5B suggests that the Antarctic data is dominated by biotechnology with pharmaceutical or medical preparations, detergents, foods, biocides and cosmetics as the other main product categories.

In terms of the number of first filings the data is clearly led by Novozymes with other companies and research organisations some distance behind. Here we would observe that Novozymes has a long standing policy of including information on the geographic origin of genetic material in patent applications. On balance, the number of filings overall and by organisation is relatively small and subject to significant yearly variation.

In practice, the emerging patent landscape for Antarctica can be divided into six main segments: a) sequence data b) *Candida antarctica*, c) Antarctic krill, d) other species recorded in the Antarctic, e) citations of the Antarctic scientific literature, f) references to Antarctic place names as collection sites. We now address each of these in turn.

### Digital Sequence Information

The prominence of biotechnology related activity is suggested by the number of Antarctic related filings containing DNA or amino acid sequences. Sequence data, under the place holder term ‘digital sequence information’ or DSI, has become an increasing focus of attention in international policy debates on access and benefit-sharing for genetic resources in recent years under the Convention on Biological Diversity and a range of other policy processes [149–154]. In the context of debates on a new treaty on marine biodiversity under the United Nations Law of the Sea, counts of genetic sequences in patent data have had a significant impact on policy debates and have attracted significant publicity [155–158].

In total 928 first filings contained a verifiable Antarctic species and DNA and amino acid sequence data. After the exclusion of records where the ubiquitous *E. coli* was the only species recorded in a document with a sequence listing, 739 first filings contained sequences.

In practice, considerable caution is required in interpreting the sequence data in patent documents. Existing research has adopted the novel approach of cumulating counts of sequences in patent documents that are linked to marine species in patent sequence data from the World Intellectual Property Organization (WIPO) [155–158]. This serves the purpose of demonstrating the increasing presence of sequences from marine organism in patent activity and links to wider questions about benefit-sharing. However, the use of cumulative counts may inadvertently disguise the reality that underlying patent filings, reflecting the outcomes of investments in research and development, may be much weaker and made up of spikes of individual documents containing large numbers of sequences [157]. Figure 8 displays three different approaches to counting sequence data.

**Figure 8:**
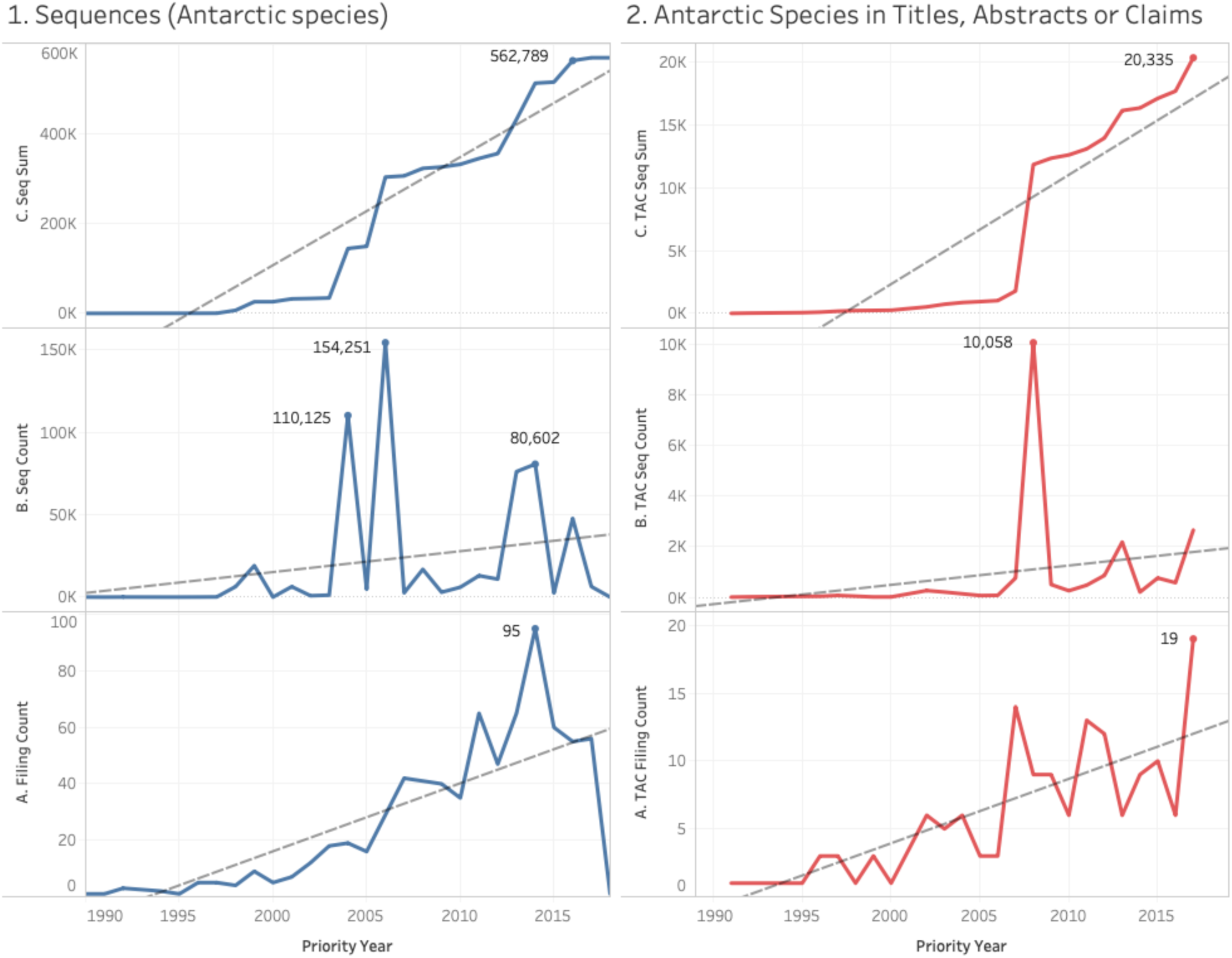
Approaches to Sequence Counts for Patent Activity for Antarctic Species.

Beginning with Figure 8(1A) we observe overall trends in first filings containing sequences for documents that also contain an Antarctic species (after the exclusion of *E. coli*). Trends in filing are clearly modest over this period and peak at 95 filings in 2014. Moving up to Figure 8(1B) we present counts of the number of sequences that appeared in documents by year. This reveals clear spikes in activity that consist of a filing in 2004 containing 108,053 sequences, a filing in 2006 containing 150,913 sequences and a set of 7 filings in 2015 containing 78,771 of 80,602 sequences recorded that year. The significance of this becomes clearer when we consider Figure 8(1C) which displays the cumulative sum over time leading to a total of 562,789 sequences. This may readily give an impression of significant commercial interest until we recognise that 46% of activity over the period is made up of two filings rising to 60% of activity across the 9 filings mentioned above. In short, cumulative trends can radically amplify otherwise weak underlying activity.

It is common practice in patent analytics to focus on documents where a subject of interest appears in the titles, abstracts or claims on the basis that the document will in a fundamental way be ‘about’ that subject. Figure 8(2) reproduces the approach in Figure 8(1) but restricts the data, after the exclusion of *E. coli*, to filings where an Antarctic species appears in the titles, abstracts of claims (TAC) of a filing. As the irregularity of this pattern in Figure 8(2A), and the associated spike in Figure 8(2B), serve to highlight, when viewed from this perspective commercial interest in Antarctic species, as reflected in sequence data, can be reasonably be described as emergent rather than intense.

A need for caution in approaching sequence data in patent filings is also reflected in the fact that, as Jefferson et. al. 2013 have ably demonstrated, sequences may appear in patent data either because they are comparative reference sequences, or because they are claimed [159]. However, disentangling referenced and claimed sequences requires close interpretation of patent claims and represents a weak area in existing methods in patent analytics. Tools such as *PatSeq* from the Lens are opening up the possibility of greater rigour in the interpretation of sequence data in patent documents.

In our view, cumulative counts of sequences can serve as a useful indicator of growing commercial interest in biodiversity in areas such as the Antarctic but should not be used in isolation from conventional counts. Cumulative counts are particularly useful for amplifying an otherwise weak signal. However, the method should logically only be used in conjunction with other counts in order to avoid giving a misleading impression of intense commercial interest in genetic resources when in practice activity is weak or emergent. Furthermore, an exclusive focus on sequence data in the case of marine genetic resources has occurred at the expense of recognition that the majority of patent activity for biodiversity and marine biodiversity does not involve sequences [157,160]. Thus, in the case of the Antarctic data presented here the 928 filings containing sequences constitute 34% of the 2,738 first filings containing an Antarctic species. As such, a broader view that accommodates the full spectrum of patent activity for biodiversity is appropriate.

#### Candida antarctica

As noted above, the type specimen for *Candida antarctica* (accepted name *Moesziomyces antarcticus*) was originally collected from sediment in Lake Vanda. Figure 9 displays an overview of filing activity for *Candida antarctica*.

**Figure 9:**
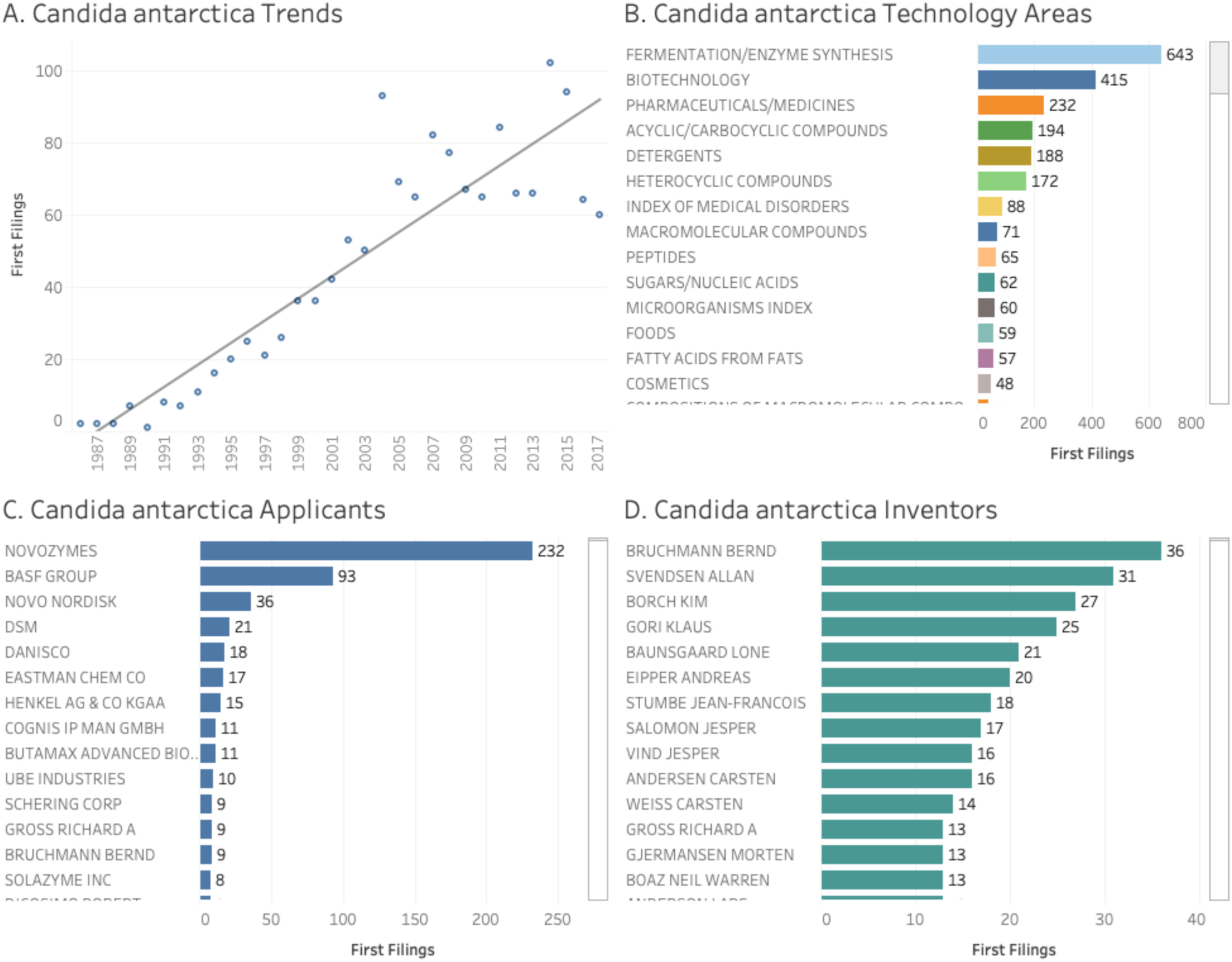
Overview of First Filings for Candida antarctica.

*Candida antarctica* is a yeast species that is a source of industrially important lipases. A lipase is any enzyme that catalyses the hydrolysis of fats. The earliest filing in the available data for *C. antarctica* can be traced to 1986 by Novozymes for the Enzymatic Synthesis of Waxes focusing on *Mucor miehei* and providing examples using *C. antarctica* linked to an earlier filing in Denmark [161]. However, the most highly cited patent document is a 1992 filing by Novo Nordisk, the original parent of Novozymes, that claims *C. antarctica* lipase and its variants including a number of modified amino acid sequences [162]. From these relatively early beginnings the use of *Candida antarctica* lipase has expanded into a variety of different sectors including medical, detergents, fuels and food stuffs for which we provide brief examples.

As we can see in Figure 9B a significant number of medical related patent applications involve *C. antarctica*. These documents typically take the form of references to the actual or potential use of the lipase in medical compositions rather than claims to the lipase itself. Thus, Neose technologies Inc claim an invention that relates to mutants of Fibroblast Growth Factor (FGF), particularly FGF-20 and FGF-2 1, which contain newly introduced N-linked or O-linked glycosylation site(s). The application also discloses polynucleotide coding sequences for the mutants, expression cassettes comprising the coding sequences and cells expressing the mutants [163]. In a similar way, Rigel Pharmaceuticals Inc disclose 2,4-pyrimidinediamine compounds having antiproliferative activity, compositions comprising the compounds and methods of using the compounds to inhibit cellular proliferation and to treat proliferate diseases such as tumorigenic cancers [164]. In our view the majority of medically focused references are likely to involve the actual or potential use of the lipase rather than direct claims involving *C. antarctica*. However, more direct use of the lipase is reflected in a University of Georgia Research Foundation Inc filing describing novel structured lipids and their use in modulating total cholesterol levels [165].

In the case of biodiesel, Wechtech Biotech Co. Ltd have applied for a method for enhancing the activity of an immobilized lipase they claim is useful in a method of preparing biodiesel by transesterification of triglycerides [166]. In the case of foodstuffs, Aker Biomarine report on novel compositions containing conjugated linoleic acids that are efficacious as animal feed additives and human dietary supplements that use *C. antarctica* lipase in the esterification process [167]. Senomyx Inc have reported that certain non-naturally occurring, non-peptide amide compounds and amide derivatives are useful flavour or taste modifiers for food, beverages, and other comestible or orally administered medicinal products or compositions [168]. However, the *C. antarctica* appears to be simply referenced in this application.

Patent activity for *C. antarctica* illustrates the point that species can be said to enjoy careers inside the patent system. These careers typically start with filings on the discovery of a useful property of an organism, are followed by claims to variants of that property and then expand to the actual or potential use of that element in a wider range of claimed inventions and products. As the uses of an element of an organism become established, research will also typically turn to identifying other useful properties of an organism and the increasing pursuit of alternatives from other sources to compete with those elements. Over time, the bulk of activity relates to the actual or potential use of the elements of an organism in a claimed invention rather than direct claims to elements of the organism. Experience suggests that the careers of many species in the patent system follow this type of pattern and this can also be observed in the case of Antarctic krill [160].

#### Antarctic krill

Antarctic krill *Euphausia superba* has become an increasing focus for the development of commercial and consumer products involving krill oil and the use of krill in feed for commercial aquaculture. Previous work by Foster et. al. 2011 highlighted the proliferation of patent activity across sectors for krill and its implications for predicting trends in krill fishery [169].

Across both scientific and common names for krill we identified 150 first filings linked to a total of 1,193 family members worldwide. We would note that this data is confined to filings that make reference to *Euphausia superba* or Antarctic krill within the Antarctic patent dataset and does not consider wider references for the simple term krill in patent documents (supplementary material). Figure 10 displays an overview of the data on first filings for Antarctic krill.

**Figure 10:**
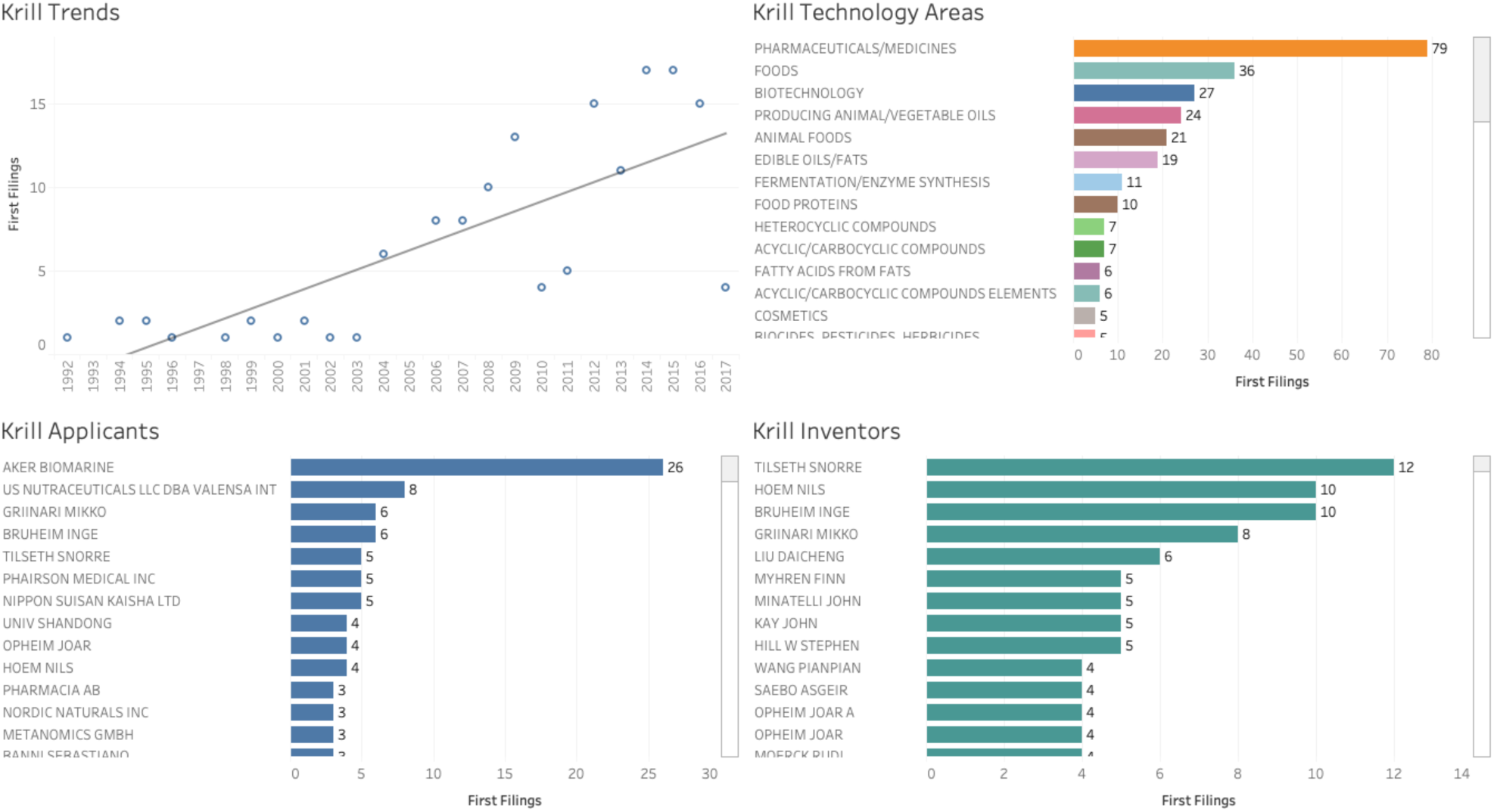
Antarctic krill.

Figure 10 reveals that while first filings in relation to krill are relatively small, there is a distinct rise in filings reflecting wider interest in commercial research and development using krill. Figure 10 focuses on the very first filings of patent applications. In contrast, Figure 11 expands the landscape to focus on all known follow on applications and grants around the world that form ‘family members’ of the first filings.

**Figure 11:**
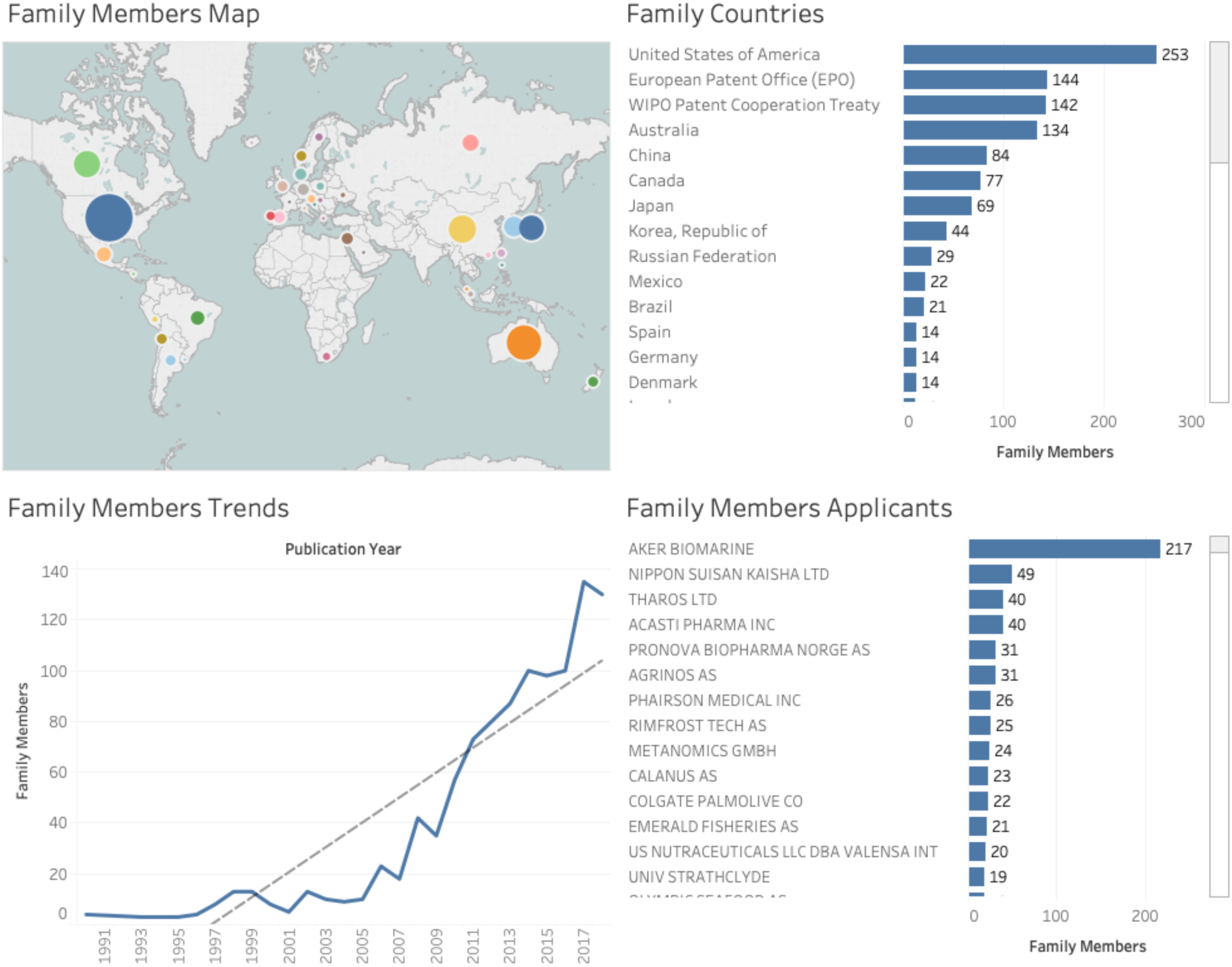
Patent Family Members Worldwide for Antarctic Krill.

Comparison between Figure 10 and Figure 11 helps to clarify that a single application may lead to multiple applications and grants around the world. Applicants must pay fees at each stage of the application procedure and, where relevant, maintenance fees for patent grants in each country. Follow on filings therefore reflect the importance of the claimed inventions to the applicants in specific markets. This data also demonstrates that a relatively small number of filings can have a wider global impact as applicants seek to protect and commercialise their claimed inventions in multiple markets. However, while Figure 11 shows a steeply rising trend the numbers are not dramatic relative to activity in the wider patent system.

In the case of Antarctic krill we are witnessing a combination of an increasing number of claims to elements of krill, such as krill oil, and the use of krill as an actual or potential ingredient in a claimed invention (such as a foodstuff, animal feed or cosmetic). In practice, filings relating to Antarctic krill can be traced back to the 1980s and the scientific literature on krill has played a significant role in promoting commercial research and development. Thus, a 1986 article on ‘Supercritical carbon dioxide extraction of oils from antarctic krill’ by researchers from Japan has been cited in the patent literature over 55 times [170]. Claimed inventions citing this article include the extraction of polar lipids and phosopholipids from krill [171], and a new krill oil composition which was found to be useful as an anti-inflammatory, as an anti-oxidant and for improving insulin resistances and blood lipid profiles [172]. We also observe activity for a krill extract aimed at treating thrombosis [173] and a method for using krill oil to treat risk factors for cardiovascular, metabolic and inflammatory disorders [174] as well as therapeutic phospholipid compositions for treating or preventing a wide range of diseases such as cardiovascular and neurodegenerative diseases [175]. The use of krill as krill meal in aquaculture has also emerged as a significant focus of commercial research and development such as krill meal products [176], as well as methods for making krill meal [177] and using krill meal as a supplement [178,179]. Recent applications include applications seeking to tackle the harmful effects of oxidised LDL cholesterol [180], and to provide nutritional supplements [181] and new lipids [182].

### Antarctic literature cited in patent documents

The prominence of *Candida antarctica*, Antarctic krill and the sheer diversity of species that appear in patent documents that mention Antarctica can make it difficult to assess activity for other Antarctic species. However, the Lens database has pioneered efforts to link the scientific literature and citing patent documents. This means that it is possible to identify and explore cases where Antarctic research is cited in a patent document.

It is important to note that a scientific publication on Antarctic biodiversity may appear in a patent document for a number of reasons. In some countries, such as the United States, applicants are required to disclose all potentially relevant prior art (scientific publications and patents) at the time of application. This can take the form of passing references that are not in reality relevant to the claimed invention. In other cases, literature on Antarctica may form part of a wider thematic set (such as anti-freeze proteins) that indirectly informs the claimed invention. In a third case, an element of an Antarctic species identified in the literature may directly form part of a composition, method or process. Finally, in a small number of cases, Antarctic researchers are both publishing and applying for patent protection for biodiversity components arising from their research. We now briefly explore this data.

The article on Antarctic biodiversity that has received the most patent citations, with over 60 citations, is a review entitled “Developments with Antarctic microorganisms: culture collections, bioactivity screening, taxonomy, PUFA production and cold-adapted enzymes” [183]. Patent applications citing this article have focussed on the production of polyunsaturated fatty acids (PUFAs) from bacterial microorganisms [184–187], including the production of the PUFA omega-3 [188]. A filing by Martek Biosciences on polyunsaturated fatty acid (PUFA) polyketide synthase (PKS) systems using *Shewanella japonica* and *Shewanella olleyana* also states that *S. olleyana* was sourced from the Australian Collection of Antarctic Microorganisms (ACAM) as strain number 644. However, the accompanying literature citation for the sample makes clear that the specific sample was from an estuary from Australia [189]. As such, while the Antarctic literature informs the claimed invention, this is an example of indirect influence.

An article on the Antarctic nematode, *Panagrolaimus davidi* that survives intracellular freezing has received 44 literature citations and is cited in 13 patent families [190]. The most highly cited patent families are from Zeltiq Aesthetics Inc and pertain to methods for cooling and treating subcutaneous lipid rich cells such as adipose tissue [191,192], and methods for interrupting or resuming treatments [193,194]. This is a second example where the Antarctic literature indirectly informs or inspires a claimed invention because the invention itself is a physical device for cooling tissue.

Patent claims involving biodiversity may be constructed on different taxonomic levels such as species, genus, family and order. In the case of order level claims, a 1974 article “Four new species of thraustochytrium from Antarctic regions…” [195] is referenced in 12 patent documents from 3 patent families filed by Martek Biosciences. However, the specific reference to Antarctica is limited to comparison with the growth conditions of other Thraustochytrium. Patent documents within the three families include a process for growing Thraustochytrium and a food product which includes Thraustochytrium [196] and processes for growing microorganisms of the order Thraustochytriales [197,198]. The first claim of one filing is for: “A process for culturing a microorganism of the order Thraustochytriales…” in a culture medium to obtain PUFA lipids. In this case it is *the process* for obtaining the lipids from the organisms that is the focus of the invention rather than biochemical compounds from the organisms *per se* as in claims for compositions of matter [196].

Examples of patent claims at the genus level are provided in a set of 18 patent applications citing an article defining the genus *Nocardiopsis*, including *Nocardiopsis antarctica*, [199]. These patent documents include direct claims relating to *Nocardiospis*, such as a filings by Novozymes in relation to proteases and associated DNA and amino acid sequences, but use species other than *N. antarctica* such as *N. alba* [200]. However, these types of application commonly anticipate the use of the same, or substantially similar sequences, from other members of the genus through reference to other species, such as *N. antarctica* elsewhere in the application.

As these examples illustrate, patent documents involving biodiversity and the biodiversity literature may inform claimed inventions in a variety of ways and require considerable care in interpretation. We now turn to patent filings that cite the Antarctic literature where an Antarctic species is directly material to the claimed invention.

An article exploring exopolysaccharides produced by marine bacteria found in Arctic and Antarctic sea ice and other extreme environments has been cited in 10 patent families [201]. These include the use of exopolysaccharides in compositions to treat subterranean formations [202] while other filings refer to the use of bacterial exopolysaccharides in cosmetic compositions, with antioxidant properties [203], anti-wrinkle properties [204], and controlling sebum secretion in the skin [205].

An article identifying the mechanisms through which Antarctic microalga *Chlorella vulgaris* is able to adapt to cold conditions and high salinity [206] has been cited in 6 patent families (10 documents). These include the use of *Chlorella vulgaris* in the production of natural oil for the purpose of manufacturing transportation fuels such as renewable diesel, biodiesel, and renewable jet fuel, as well as oleochemicals such as functional fluids, surfactants, soaps and lubricants [207]. This patent application has been cited by over 17 later filings. Another patent application utilising the species in the production of renewable fuels, which are also useful as feedstocks, also cites this article [208].

Research on alkaloids from the Antarctic sponge *Kirkpatrickia varialosa* in the mid 1990s has been cited in four patent families containing 10 documents led by the Spanish pharmaceutical and marine biodiscovery company Pharma Mar filed from 2000 onwards [209]. The patent families focus on the anti-tumour properties of Variolin and its derivatives [210–213]. Three of the patent families contain over 30 family members with protection sought in 21 countries suggesting that the applicants believe that the claimed invention has significant commercial potential.

As discussed above, cold tolerance or antifreeze molecules and proteins have been a significant area of research in the Antarctic. The Antarctic grass *Deschampsia antarctica* has been a significant focus of Antarctic research with 251 articles in our Antarctic literature dataset with top cited scientific literature focusing on issues such as heat tolerance of photosynthesis, the evolution of UV absorbing compounds, and vascular plants as bioindicators for warming in Antarctica [214–216].

Scientific literature that is cited by patent applicants includes work on three cold-responsive genes from *Deschampsia antarctica* by researchers from Chile [217]. This research is cited by three patent families including one for an ice recrystallisation inhibition protein, and another for an isolated low temperature plant promoter gene [218,219]. Other work of relevance includes work on the characterization of antifreeze activity in Antarctic plants [220] that is cited in a 2013 patent grant for an agent for cutaneous photoprotection against UVA (I and II) and UVB radiation (skin protection against sun damage) containing an aqueous extract from *Deschampsia antarctica* either obtained from its native environment or grown in artificial settings [221]. As this suggests, genetic elements and compounds from Antarctic species may find applications in multiple industry sectors. In total, as highlighted in Figure 5, we identified 26 first filings involving *Deschampsia antarctica*.

An article examining the antifreeze protein gene from the antarctic marine diatom *Chaetoceros neogracile* [222] is cited in a 2014 patent family filed by Samsung electronics for an “Antifreeze Member”. The focus of the claimed invention is the creation of a metal substrate for semiconductors, energy and biosensors that overcomes the problem of frost formation on cooling plates. A 2017 US patent grant to Samsung claims that this problem can be solved by “a recombinant antifreeze protein in which a metal-binding protein is conjugated to an antifreeze protein derived from Chaetoceros neogracile” [223].

In what appears to be a small number of cases the authors of scientific articles are also applying for patent protection. One example is work by researchers in Korea from the Korea Polar Research Institute and the Korea Ocean Research and Development Institute in work on the antioxidant properties of lichens from Antarctica, notably *Ramalina terebrata* [224]. In this case the research has led to the filing of 5 applications focusing on Ramalin from *Ramalina terebrata* [225,226]. This includes the use of Ramalin for its antioxidant properties, in pharmaceutical products to treat oxidation related diseases, in functional foods for anti-aging purposes and in functional cosmetics for skin-whitening and anti-wrinkle purposes [227]. One patent application relates to the use of Ramalin in a pharmaceutical composition to treat or prevent inflammatory and immune diseases [228]. Another application relates to anti-cancer treatment for colorectal cancer [229]. Taken together the filings suggest a strategy to capture a broad range of potential medical applications for Ramalin. At the time of writing the scientific landscape for Ramalin consists of 31 scientific publications and 21 patent families. This represents a significant research investment and strongly suggests that the applicants believe Ramalin has commercial potential.

Corals and tunicates, such as sea squirts, have been a major focus of applied and commercial marine research [230]. In the case of Antarctica *Synoicum adareanum* has been the subject of research on a cytotoxic macrolide that also formed the basis for a patent application and grant to the lead authors [231,232]. The sea squirt *Aplidium cyaneum* has also been a focus of research on cytotoxic bromoindole derivatives that became the basis of a patent application by some of the authors [233,234].

Antarctic fish have also become a significant focus of commercially oriented research and development. The scientific literature has focused on issues such as the role of Notothenioid fish in the food web of the Ross Sea shelf [235], or neutral buoyancy in Notothenioid [236]. Commercially oriented research for Notothenioids such as *Dissostichus mawsoni* focuses on antifreeze glycopeptides in the tissues and fluids of Antarctic fish [237] and comparative analysis of these proteins between Arctic and Antarctic fish [238]. This work has resulted in a direct filing in 1990 by at least one of the researchers at the University of California for thermal hysteresis proteins with a significant impact on later patent filings in the form of 57 patent citations focusing on issues such as ice-controlling molecules and cryosurgery [239]. In total 7 first filings relating to *Dissostichus mawsoni* were identified in the patent dataset.

Other Notothenioidei that are a focus of commercial research and development include the White Blooded Icefish (*Chaenocephalus aceratus*) [240,241]. Work on icefish lacking in haemoglobin is reflected in a 1999 filing on methods for the isolation of hemapoietic genes in Antarctic icefish [242]. Comparative research involving *Chaenocephalus aceratus* focusing on Vitamin E content [243] associated with cold adaptation has also attracted a patent citation but with a specific focus on a krill composition [244]. *Pagothenia borchgrevinki* is also a source for a patent filing by Airbus in 2008 for anti-freeze proteins for application to wings, rotors and turbines [160,245].

As these examples make clear, analysis of patent documents that cite the Antarctic literature provide a clear route to monitoring filings where an Antarctic species is material to a claimed invention. However, care is required in interpreting the reasons why an article is cited and whether an Antarctic species is directly involved or material to the claimed invention. We conclude this exploration of the patent landscape by briefly examining references to Antarctic place names in patent data.

### Antarctic Places

One aim of the research was, as far as possible, to identify and map place names appearing in the literature and patent data. Two main sources of data are available for places in Antarctica. The first is the SCAR Composite Gazetteer (March 2018) of 36,630 names. The second source is the Geonames database, which produces a file AQ for Antarctica containing 18,526 place names and 27,273 variant names in multiple languages. To examine references to Antarctic places in patent documents we deduplicated the names and then mapped the roots of place names into the patent data focusing on patent documents that also contain a species name. In total we identified 267 filings that contained a reference to a place name and a species name, dominated by the term Antarctic/Antarctica. References range from general descriptions of krill as an Antarctic species to Antarctic islands. Here we focus on illustrative examples.

A 2010 filing by researchers from the Korea Ocean Research Development Institute (published as EP2617464A1) makes multiple references to places including King Sejon Station, Barton Peninsula and King George Island. The application focuses on Antarctic lichens notably an extract of *Stereocaulon alpinum* in pharmaceutical and food compositions to prevent or treat diabetes or obesity and has a patent family with 15 members including patent grants in China, under the European Patent Convention, Japan and the United States [246]. The patent application and other members of the patent family explain that:

“…the Antarctic lichen Stereocaulon alpinum (Stereocaulon alpinum (Hedw.) G.L. Sm.) used in the present invention was collected from the area around the King Sejong Station (S 62°13.3’, W58°47.0’) located on Barton Peninsula on King George Island, Antarctica, in January 2003.”

An important feature of this explicit reference is that it is possible to identify the precise point of collection through the use of a named place and coordinates. This is also a case where at least one of the authors of research on *Stereocaulon alpinum* is listed as an inventor [247].

A second example of direct collection of samples in Antarctica also reveals the close relationship between the publication of scientific articles and patent filings. A 2016 filing from researchers from the University of South Florida and the University of Alabama (UAB) Research Foundation addresses MRSA Biofilm Inhibition [248]. The application states that:

“In the course of acquiring biodiversity to support an antibiotic screening program, the current inventors obtained the sponge Dendrilla membranosa from the vicinity of Palmer Station, Antarctica. The dichlorom ethane extract of the freeze-dried sponge was subjected to reversed-phase solid-phase extraction eluted with acetonitrile. The extract underwent HPLC purification to yield four major natural products, including three previously reported spongian diterpenes: aplysulphurin, tetrahydroaplysulphurin, and membranolide (Karuso et al., Aust. J. Chem. 1984, 37, 1081-1093; Karuso et al., Aust. J. Chem. 1986, 39, 1643-1653; and Molinski et al., J. Org. Chem. 1989, 54, 3902-3907). The fourth product was identified as darwinolide, a new rearranged spongian diterpene having a structure shown in FIG. 1. … The darwinolide skeleton is the newest of over a dozen structural motifs distinguishing the broad chemodiversity found in the Darwinellidae family of sponges.” [248]

They go on to explain that:

“Sponge samples were collected from various sites around Palmer Station, Antarctica in the austral summer of 2011. The collection sites chosen were Norsel Point (64°45.674’ S, 64°05.467’W), Bonaparte Point (64°46.748’ S, 64°02.542’W), Gamage Point (64°46.345’ S 64°02.915’W), and Laggard Island (64°48.568’ S, 64 00.984’W) at depths between 5-35 m below sea level. Samples were frozen and transported back to the University of South Florida at -70°C where tissues were lyophilized and stored at - 80°C until further processing.”

The applicants claim a method for treating bacterial infections including MRSA biofilms with a darwinolide compound. However, this is also a case where researchers time the submission of a scientific article and a patent filing in such a way that the research article, which would become prior art, does not destroy the novelty of the claimed invention. Thus, the earliest filing date of the patent application is in April 2016 shortly before the publication of the scientific article in May 2016 and is followed in October 2017 by publication of the Patent Cooperation Treaty patent application [249].

A third example highlights that applicants may obtain samples through Antarctic research centres operating as intermediaries. A 2009 first filing from India became the basis for a 2010 international Patent Cooperation Treaty application [250] for methods of preparing a plant extract using liquid chromatography and mass spectrometry where the plant extract is from *Deschampsia antarctica*. This application describes how:

“The frozen plant material was procured from Coppermine Peninsula on Robert Island, South Shetland Island, Antarctica and was exported to us by Instituto Antarctico Chileno…”

This application also makes extensive reference to the wider literature on *D. antarctica* that mention places such as Signy Island and King George Island signifying that the intensity of occurrences of references to Antarctic places may be a good indicator of collection of samples in the Antarctic. However, the main insight from this example is that in some cases an Antarctic research institute may serve as an intermediary providing Antarctic material for commercially oriented research. It is unclear whether the institute was aware of this purpose when providing the material or whether a material transfer agreement (MTA) was established between the institute and the applicants.

A fourth example illustrates the point raised above that a sample may come from multiple sources. A 2006 filing by the Monterey Bay Aquarium Research Institute for “A light-driven energy generation system using proteorhodopsin” explains that:

Using the same proteorhodopsin-specific PCR primers, as for instance shown in FIGS. 2 and 3, proteorhodopsin genes were also amplified from bacterioplankton extracts. As mentioned above, any proteorhodopsin-specific PCR primer can be used. These bacterioplankton extracts include those from the Monterey Bay (referred to as MB clones), the Southern Ocean (Palmer Station, referred to as PAL clones), and waters of the central North Pacific Ocean (Hawaii Ocean Time series station, referred to as HOT clones).

A similar multi-source case is provided by a filing from Woods Hole Oceanographic Institute for metagenomic samples collected by drilling through sea ice in the Ross Sea combined with analysis of other diatoms to create a recombinant organism for the expression of Cobalamin (vitamin B12) [251]. An additional example is a filling for a cryoprotective agent from a novel *Pseudoalteromonas sp.* strain CY 01 (KCTC 12867BP) collected from the Antarctic Ocean as well as Arctic strains [252]. While these applications explicitly involve samples from the Antarctic, it can be challenging to determine whether the organisms are material to (part of) the claimed invention.

References to Antarctic place names occur in the context of wider international debates on disclosure of the origin of genetic material in patent applications and the consequences of such disclosure [253]. Increasingly, countries that are party to the Convention on Biological Diversity and its Nagoya Protocol are requiring disclosure of origin in support of the implementation of these agreements. However, the consequences of disclosure, and failure to disclose, may vary considerably. The present research reveals that applicants will often mention Antarctic origin and may, as we have just seen, be explicit about the places and coordinates of collection. In international policy debates at WIPO, agreement on international requirements for disclosure of origin has become stuck on disagreements about the consequences of disclosure, such as revocation of a granted patent in the absence of evidence of prior informed consent and a benefit-sharing agreement with the country of origin [253,254]. However, in the case of the Antarctic, as for marine biodiversity in Areas Beyond National Jurisdiction, the function of disclosure could perhaps better be seen as making the contribution of Antarctic biodiversity to innovation (as partly reflected in the patent system) visible to the wider world [157,160]. That is, disclosure can assist with supporting greater awareness of the ecosystem services provided by Antarctic biodiversity and thus of Antarctica to wider human welfare. However, debates on disclosure of origin also raise harder questions about the contribution that those who seek to commercially develop and use Antarctic biodiversity should make to its conservation. We turn to how this issue might be addressed in closing.

## Conclusion

This paper has sought to contribute to mapping the scientific and patent landscape for biodiversity and innovation in Antarctica and the Southern Ocean. The growing availability of open access databases of scientific, patent and taxonomic data means that it is possible to begin to map these landscapes at scale using methods that are open, transparent and accessible to a range of disciplines. However, as we have also sought to demonstrate, exploiting opportunities for analysis at scale reveals issues around data completeness and data quality. In the case of patent data, these challenges extend to requirements for considerable care in interpretation of the Antarctic origin of genetic resources within patent documents and whether they are actually material to or part of the claimed invention.

Issues around data completeness and data quality can be addressed through approaches such as re-indexing to address gaps, in the case of Microsoft Academic Graph, and closer attention to data cleaning using locality information for taxonomic data from GBIF. The growing availability of the full texts of both scientific and patent publications presents important opportunities to improve access to the full results of scientific research but also presents challenges in moving beyond pure metadata based approaches. Open access databases such as the Lens have made important breakthroughs by linking together scientific and patent data through citations. This in turn makes it easier to monitor patent activity arising from research involving Antarctic biodiversity. Developments in machine learning, in the form of Natural Language Processing libraries such as spaCy, mean that it is now possible to imagine a pipeline approach to monitoring Antarctic research by streaming new scientific publications and patent data from database application programming interfaces (APIs), such as the Lens, through a machine learning model for classification, name entity recognition, analysis and distribution to the scientific and policy community. The growing popularity of pipeline approaches to dealing with data at scale reflects the widespread availability of open source libraries for analytics at scale. Implementing such a pipeline would require focused investment by one or more members of the Antarctic Treaty System and would logically be coordinated with the SCAR. As this paper helps to demonstrate, this is an achievable goal.

The present research also points to potential ways forward in addressing harder questions around benefit-sharing from commercial research and development involving Antarctic biodiversity. Bioprospecting has been on the agenda of the Antarctic Treaty System for a number of years. However, as far as we are aware, beyond agreement to keep discussing the issue, no consensus has emerged on a need for practical action other than collecting more information to inform deliberations. This has a certain logic in light of uncertainties about levels of activity and the actual or potential overlap between genetic resources inside the Antarctic Treaty System, those within national jurisdictions and those being considered by debates on the new treaty on marine biodiversity in areas beyond national jurisdiction under the Law of the Sea.

One challenge with the treatment of bioprospecting, or commercial research and development as we prefer, is that it is largely seen in isolation from other activities in Antarctica. A way forward could potentially be found by viewing commercial research and development from an ecosystem services and natural capital accounting perspective. Many, if not all members of the ATS, have embraced the ecosystem services approach and a growing number are moving towards testing or implementing natural capital accounting in accordance with the framework of the System of Environmental Economic Accounting (SEEA) linked to the United Nations Systems of National Accounts (SEA) [19,20,255,256]. These developments have been accompanied by the increasing promotion of the concept of Payments for Ecosystem Services (PES) within the environmental economics literature and policy, as proposed by Verbitsky 2018 for tourism in Antarctica [19,257]. This type of approach would allow countries to draw on existing experience with ecosystem services and natural capital accounting when addressing commercial activity in the Antarctic. It should be emphasised that the valuation of ecosystem services is challenging and it is increasingly recognised that there is a risk that such approaches may seek to reduce biodiversity to an equivalent monetary value at the expense of recognition of the multiple values of biodiversity and its services. Nevertheless, despite these reservations, over the short and medium term this approach would place the assessment of activities such as commercial research and development or tourism within a clear and transparent framework that would bring Antarctica into the fold of wider work on the economics of biodiversity.

The year 2020 has been described as a super year for biodiversity. As countries scramble to address the formidable damage caused by Covid-19 it remains to be seen whether this will become a reality. However, one important lesson from the environmental and ecological economics literature is that biodiversity cannot be treated as a free good. The joint biodiversity and climate crisis has its origins in the treatment of the environment as a free good when in fact the costs are deferred elsewhere including to future generations. When viewed from this perspective, biodiversity is not free but has to be paid for. At present, as far as we are aware, the revenue generated by biodiversity based innovation from research in the Antarctic does not contribute to the conservation of biodiversity in the Antarctic. 2020 provides an opportunity to rethink the logic that produces this situation by recognising that biodiversity must be paid for. By accepting that biodiversity is not free we are then able to ask other questions focusing on returning tangible benefits to Antarctic biodiversity such as: how much, by whom, in what form, and to what ends? This paper seeks to contribute to the development of the evidence base for addressing these questions.

## Footnotes

2 Available through the Biospolar Antarctic Literature and Patents repository at https://osf.io/py6ve/

3 Publicly accessible at: https://www.lens.org/lens/collection/179814

4 Because a species name may appear in multiple parts of the same document the overall counts will be higher than the totals in Figure 5.

i https://www.microsoft.com/en-us/research/project/microsoft-academic-graph/

