## supplementary material for "Biodiversity Research and Innovation in Antarctica and the Southern Ocean"

Open access datasets are accessible through the Open Science Repository at: https://osf.io/py6ve/

A set of public collections are available on the Lens database containing the records discussed in the article. These can also be accessed through the OSF repository wiki at: https://osf.io/py6ve/wiki/home/

Literature

- Antarctic Biodiversity Literature Collection with 25,468 publications.
- Antarctic Bioprospecting Literature with 98 publications.
- Antarctic Biodiversity Literature with Patent Citations with 796 publications.

Patent Data

- Raw patent data on Antarctica, 29,690 documents
- Antarctic Species, 2,738 documents. Be aware that this set will include incidences of E. coli as discussed in the paper where it co-occurs with references to Antarctica).
- Krill first filings. The 150 first filings examined in the paper, with 1,0193 family members. Refers to filings mentioning Antarctica or Euphausia superba inside the Antarctic dataset. Note that this dataset is dynamic and expansion of patent family members may reveal more documents than listed in the paper.
- Krill. An expanded patent dataset containing patent families containing the term simple term krill (4,106 families).
- Antarctic species and places. A collection consisting of 267 raw families (not reduced to first filings) and an Antarctic place name. Dominated by references to Antarctic/Antarctica.
